# Acetylcholine Promotes Directionally Biased Glutamatergic Retinal Waves

**DOI:** 10.1101/2023.11.10.566639

**Authors:** Kathy Zhang, Ashley Su, Yixiang Wang, Michael Crair

## Abstract

Spontaneous retinal waves are a critical driving force for the self-organization of the mouse visual system prior to eye-opening. Classically characterized as taking place in three distinct stages defined by their primary excitatory drive, Stage II waves during the first postnatal week are propagated through the volume transmission of acetylcholine while Stage III retinal waves during the second postnatal week depend on glutamatergic transmission from bipolar cells. However, both late Stage II and early Stage III retinal waves share a defining propagation bias toward the temporal-to-nasal direction despite developmental changes in the underlying cholinergic and glutamatergic retinal networks. Here, we leverage genetic and pharmacological manipulations to investigate the relationship between cholinergic and glutamatergic neurotransmission during the transition between Stage II and Stage III waves *in vivo.* We find that the cholinergic network continues to play a vital role in the propagation of waves during Stage III after the primary mode of neurotransmission changes to glutamate. In the absence of glutamatergic waves, compensatory cholinergic activity persists but lacks the propagation bias typically observed in Stage III waves. In the absence of cholinergic waves, gap junction-mediated activity typically associated with Stage I waves persists throughout the developmental window in which Stage III waves usually emerge and lacks the spatiotemporal profile of normal Stage III waves, including a temporal-to-nasal propagation bias. Finally, we show that cholinergic signaling through β2 subunit-containing nicotinic acetylcholine receptors, essential for Stage II wave propagation, is also critical for Stage III wave directionality.

## Introduction

In the absence of external sensory drive, the developing mouse retina spontaneously generates propagating waves to drive the self-organized functional maturation of the visual system (Ackman & Crair, 2014; Blankenship & Feller, 2010). By providing topographically correlated activity between presynaptic retinal ganglion cells (RGCs) and postsynaptic targets, spontaneous retinal waves have been implicated in the maturation and maintenance of retinotopic and eye-specific projections from the retina to the superior colliculus (SC) and dorsal lateral geniculate nucleus (dLGN) of the thalamus (Dhande et al., 2011; Xu et al., 2011). Beyond gross circuit formation, specific spatiotemporal characteristics of retinal waves also play a role in the development of visual neuron response properties in downstream SC neurons (Ge et al., 2021; Wang et al., 2009).

Retinal waves progress in three chronological stages (Stages I, II and III) defined by the underlying circuit driving their propagation. Stage I waves beginning around embryonic day 16 (E16) depend on spontaneous activity generated by RGCs propagating through gap junctions (Blankenship & Feller, 2010; Voufo et al., 2023). In the first postnatal week, Stage II waves propagate through the volume transmission of acetylcholine from starburst amacrine cells (SACs) to downstream RGCs through β2 subunit-containing nicotinic acetylcholine receptors (β2-nAChRs) (Bansal et al., 2000; Ford & Feller, 2012). Stage III waves during the second postnatal week persist until eye-opening, around postnatal day 14 (P14), and depend on the spontaneous release of glutamate from bipolar cells (BCs) onto RGCs through ionotropic glutamate receptors (Blankenship et al., 2009; Blankenship & Feller, 2010).

Previous *in vitro* work has provided insights into the circuit mechanisms underlying the transitions between different stages of retinal waves. In the absence of cholinergic waves, achieved by using a genetic knockout of the β2-nAChRs, residual spontaneous activity mediated by gap junctions persists and leads to the precocious emergence of glutamatergic activity (Bansal et al., 2000; Stacy et al., 2005; Sun et al., 2008). In the absence of glutamatergic waves, achieved by using a genetic knockout of the vesicular glutamate transporter 1 (VGLUT1), persistent spontaneous activity is mediated by continued cholinergic transmission (Blankenship et al., 2009). These studies suggest that the cessation of each mode of transmission is facilitated by the onset of the next mode of transmission. However, whether the excitatory network facilitating the preceding stage of retinal waves plays a continued role in the maintenance of the following stage of retinal waves remains unclear. For example, previous work has shown that pharmacological inhibition of acetylcholine severely disrupts Stage III glutamatergic waves in the ferret, but not the mouse (Bansal et al., 2000; Davis et al., 2015; Sun et al., 2008).

During the transition from Stage II to Stage III waves, BCs become integrated into the retinal circuit and SACs undergo physiological and morphological changes that cause them to become more electrically isolated and less excitable, thus losing the ability to propagate retinal waves (Ford & Feller, 2012). Simultaneously, SACs develop the necessary cholinergic and GABAergic synapses onto post-synaptic direction-selective ganglion cells (DSGCs) to mediate their direction-selective response properties (Wei et al, 2011). Whether SACs and cholinergic transmission continue to participate in Stage III glutamatergic retinal waves is unclear, but SAC expression of glutamate receptors at this age suggests a possible role (Hall et al., 2019; Hellmer et al., 2021; Shekhar et al., 2016). Despite developmental changes in the underlying retinal circuitry, both late Stage II and emerging Stage III waves share a propagation bias from the temporal-to-nasal direction in the retina that is critical for the development of direction-selective responses in downstream SC neurons and requires asymmetric GABAergic inhibition provided by SACs (Ge et al., 2021). Whether cholinergic input from SACs might play a role in mediating glutamatergic wave directionality has not been previously studied.

Here, we investigate the relationship between cholinergic and glutamatergic networks during the transition between Stage II and Stage III waves *in vivo* and demonstrate that cholinergic Stage II waves are necessary for the emergence of glutamatergic Stage III retinal waves. Further, we find that acetylcholine through β2-nAChRs is necessary for biased wave propagation during Stage III waves and propose a circuit model for Stage III wave directionality.

## Results

### In the absence of glutamatergic waves, abnormal cholinergic activity persists *in vivo* at P9-11

Stage III retinal waves, which typically begin around P10 and end shortly before eye-opening, are facilitated by spontaneous glutamate release provided by BCs known to exclusively express vesicular glutamate transporter 1 (VGLUT1) (Fremeau et al, 2004; Wässle et al., 2006). Previous experiments performed *in vitro* have shown that mice which lack VGLUT1 exhibit spontaneous activity at P10-12 that is blocked by DHβE, a nicotinic acetylcholine receptor antagonist, but not by a combination of DNQX or NBQX and AP5, ionotropic glutamate receptor antagonists (Blankenship et al., 2009). These findings suggest that continued cholinergic transmission sustains spontaneous activity in the absence of Stage III glutamatergic retinal waves.

To test whether cholinergic activity persists in the absence of glutamatergic waves *in vivo*, we utilized a mutant mouse line (*Vsx2-SE* gene KO mice) that fails to develop BCs (Gamlin et al., 2020; Norrie et al., 2019). We performed intraocular injections of AAV2/9-hSyn-GCaMP6s virus at P0-1 to label RGCs for calcium imaging of retinal axons projecting to the SC. By crossing Vsx2-SE KO mice with Emx1-Cre; Tra2β^f/f^ mice that develop without a cortex, we were able to image the full extent of the SC and thus observe the full extent of retinal activity due to the topographic preservation of projections from the retina to SC (Figure 1A) (Shanks et al., 2016). We performed widefield one-photon calcium imaging at P9-11 of spontaneous retinal activity in head-fixed, awake homozygous Vsx2-SE KO mice (Vsx2-SE^−/−^) and littermate heterozygous controls (Vsx2-SE^−/+^).

**Fig. 1.**
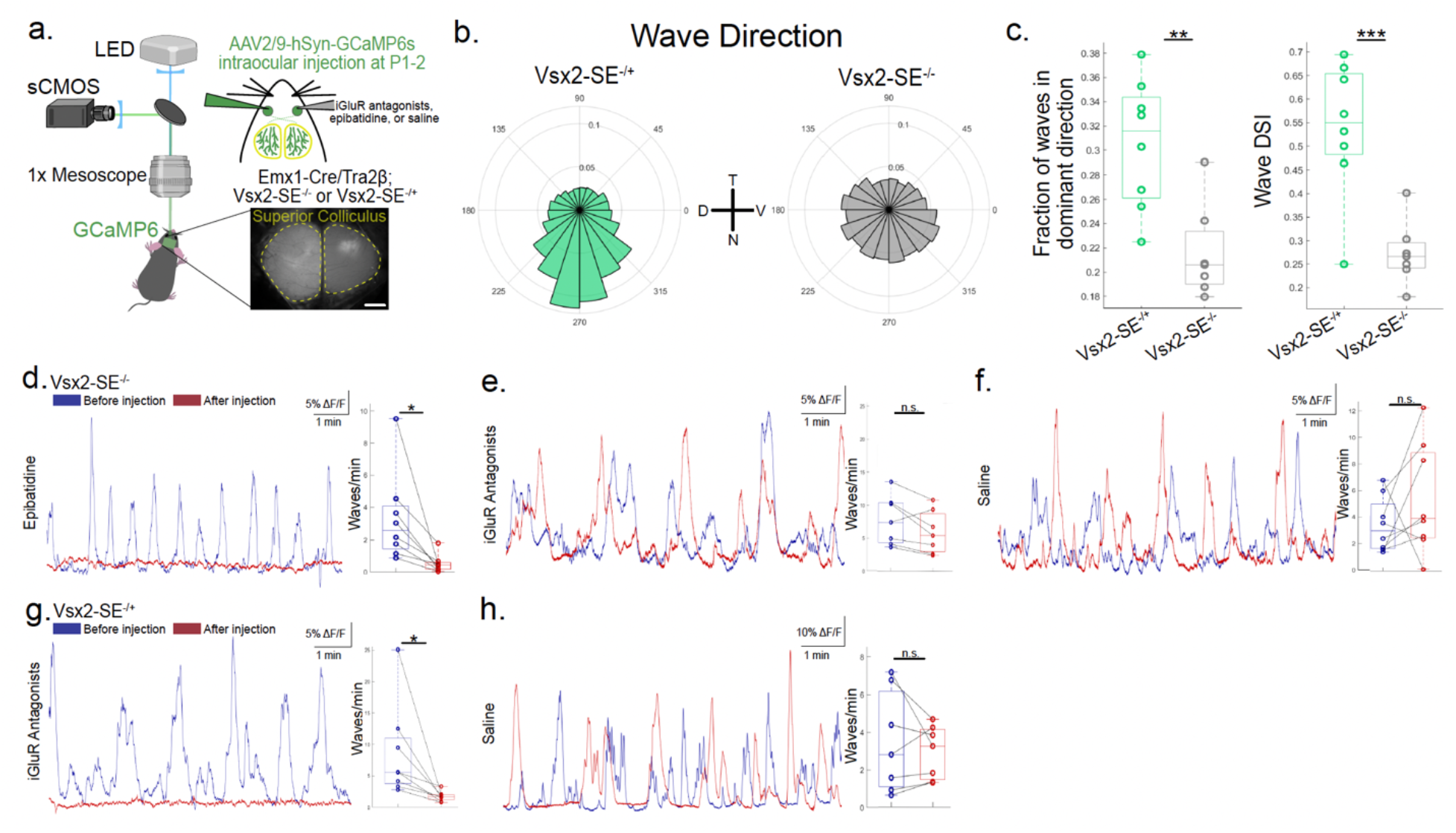
In the absence of glutamatergic waves, abnormal cholinergic activity persists *in vivo* at P9-11. **a**. Schematic of widefield single-photon microscopy used to perform calcium imaging of retinal ganglion cell axon activity in the superior colliculus (SC). GCaMP6s expression in retinal ganglion cells was achieved by intraocular injections of AAV2/9-hSyn-GCaMP6 at P1 or P2 in Emx1-Cre/Tra2β^f/f^ (cortex-less mice) crossed with Vsx2-SE^−/−^ or Vsx2-SE^−/+^ mice. A cranial window was surgically prepared before imaging at P9-11 (see Methods for details). Mice were unanesthetized and head-fixed during imaging. Spontaneous activity was recorded before and after an acute intraocular injection of either a cocktail of ionotropic glutamate receptor (iGluR) antagonists, epibatidine, or saline. Scale bar, 500 μm. **b**. Frequency distribution of propagation directions of waves in Vsx2-SE^−/+^ (n = 8 animals) and Vsx2-SE^−/−^ mice (n = 7 animals). Inset: Directions T, N, D, and V correspond to the temporal, nasal, dorsal, and ventral directions in the retina. **c**. Left: Proportion of waves traveling within 60° around the dominant temporal-to-nasal direction. Each data point represents one animal; p = 0.0026. Unpaired t-test. Right: Wave DSI is calculated as the difference between the proportion of waves traveling within 60° around the dominant temporal-to-nasal direction and the proportion of waves traveling within 60° around the opposite nasal-to-temporal direction, divided by the sum. Each data point represents one animal; p = 6.03e^−4^. Unpaired t-test. **d**. Left: ι1F/F traces of calcium transients averaged across all pixels from one hemisphere of the SC before (navy) and after (maroon) an acute injection of 100 μM epibatidine in Vsx2-SE^−/−^ mice. Right: Number of waves detected per minute before and after the injection. Each data point represents one animal, lines connect data from the same animal. n = 8 animals; p = 0.02. Paired t-test. **e**. Left: ι1F/F traces of calcium transients before and after an acute injection of a combination of 1 mM CP 465022 and 5 mM APV in Vsx2-SE^−/−^ mice. n = 7 animals; p = 0.47. Paired t-test. **f**. Left: ι1F/F traces of calcium transients before and after an acute injection of saline in Vsx2-SE^−/−^ mice. n = 8 animals; p = 0.29. Paired t-test. **g**. Left: ι1F/F traces of calcium transients before and after an acute injection of a combination of 1 mM CP 465022 and 5 mM APV in Vsx2-SE^−/+^ mice. n = 7 animals; p = 0.03. Paired t-test. **h**. Left: ι1F/F traces of calcium transients before and after an acute injection of saline in Vsx2-SE^−/+^ mice. n = 7 animals; p = 0.07. Paired t-test.

Despite the absence of BCs and thus glutamatergic retinal waves, we observed persistent spontaneous activity in Vsx2-SE^−/−^ mice at P9-11. However, the residual spontaneous activity lacked some of the typical spatiotemporal characteristics of retinal waves normally observed at this age. Strikingly, the residual activity in Vsx2-SE^−/−^ mice no longer exhibited a temporal-to-nasal directional bias typical of Stage III retinal waves and which was observed in littermate controls (Figure 1B) (Ge et al., 2021). The propagation bias was quantified by measuring either the fraction of waves propagating in the dominant temporal-to-nasal direction or a “Wave Direction Selective Index” (Wave DSI), calculated by taking the difference from the fraction of waves propagating in the dominant direction and the fraction propagating in the opposite direction, divided by the sum. Using either measure, Vsx2-SE^−/−^ mice exhibited a significantly reduced propagation bias compared to littermate controls (Figure 1C).

The similarity of wave directionality between Vsx2-SE^−/+^ animals and wildtype (WT) littermate controls at P9-11 and between Vsx2-SE^−/−^ and Vsx2-SE^−/+^ mice at P5-7 confirmed that Vsx2-SE^−/+^ animals were an appropriate control for typical Stage III wave directionality and that the observed abnormal wave directionality in the residual activity of Vsx2-SE^−/−^ mice at P9-11 was not due to off-target developmental effects of the mutation, respectively (Supplementary Fig 2-3).

To test whether residual spontaneous activity in Vsx2-SE^−/−^ mice was mediated by acetylcholine or glutamate, we performed intraocular injections of either epibatidine, an antagonist of nAChR-mediated activity, or a combination of CP 465022 and AP5, ionotropic glutamate receptor (iGluR) antagonists targeting AMPA and NMDA receptors, respectively (Cang et al., 2005; Corrie et al., 2020; Davis et al., 2015; Huberman et al., 2002; Lazzaro et al., 2002; Marks et al., 1996; Penn et al., 1998; Pfeiffenberger et al., 2005; Spang et al., 2000; Sun et al., 2008). We found that residual activity in Vsx2-SE^−/−^ mice was blocked by epibatidine but not by iGluR antagonists, suggesting that activity in the absence of Stage III glutamatergic waves is mediated by sustained cholinergic activity as has been previously reported *in vitro* (Video 1 and Figure 1 D-E) (Blankenship et al., 2009). The same combination of iGluR antagonists significantly attenuated spontaneous retinal waves in control Vsx2-SE^−/+^ animals while saline had no significant effect in either Vsx2-SE^−/−^ or Vsx2-SE^−/+^ mice (Video 2 and Figure 1F-H).

### In the absence of cholinergic waves, Stage III glutamatergic retinal waves fail to emerge *in vivo* at P9-11

The persistence of cholinergic activity in the absence of Stage III glutamatergic retinal waves corroborates previous *in vitro* findings and supports the hypothesis that the emergence of the glutamatergic network facilitates the disassembly of the cholinergic network and thus the cessation of cholinergic retinal waves. However, whether the reverse might be true, that the presence of the cholinergic network is necessary for the emergence of glutamatergic retinal waves, remains unclear. Previous work has shown that in β2-nAChR KO mice which lack cholinergic waves, gap junction-mediated activity characteristic of Stage I waves persists between P0-7 followed by the premature emergence of Stage III glutamatergic waves around P8 (Burbridge et al., 2014; Stacy et al., 2005; Sun et al., 2008; Bansal et al., 2000). These findings suggest that the presence of the cholinergic network underlying Stage II retinal waves is not required for the emergence of Stage III glutamatergic waves. Here, we use the mutant β2-nAChR KO mouse line to investigate whether Stage III glutamatergic waves emerge in the absence of Stage II cholinergic waves *in vivo*.

We performed *in vivo* calcium imaging of the retinal axons projecting to the superior colliculus in head-fixed, awake mutant β2-nAChR KO mice crossed with Emx1-Cre; Tra2β^f/f^ (β2^−/−^) and their heterozygous littermate controls (β2^−/+^) at P9-11 (Figure 2A). As expected, we found that intraocular injections of iGluR antagonists significantly decreased the frequency of spontaneous retinal waves in control P9-11 β2^−/+^ mice (Video 3 and Figure 2B-D). On the other hand, in β2^−/−^ mice lacking cholinergic waves, residual spontaneous activity at P9-11 was significantly attenuated by intraocular injections of MFA, a gap junction antagonist, but not iGluR antagonists or epibatidine, suggesting that glutamate-mediated activity fails to emerge in these mice at P9-11 *in vivo* (Video 4, Figure 2E-G). These results build upon previous findings that in the absence of cholinergic waves, gap junction-mediated activity persists between P0-7 when cholinergic waves typically occur, and further suggests that gap junction-mediated activity continues through P9-11 when the emergence of glutamate-mediated activity would be expected (Burbridge et al., 2014; Stacy et al., 2005; Sun et al., 2008).

**Fig. 2.**
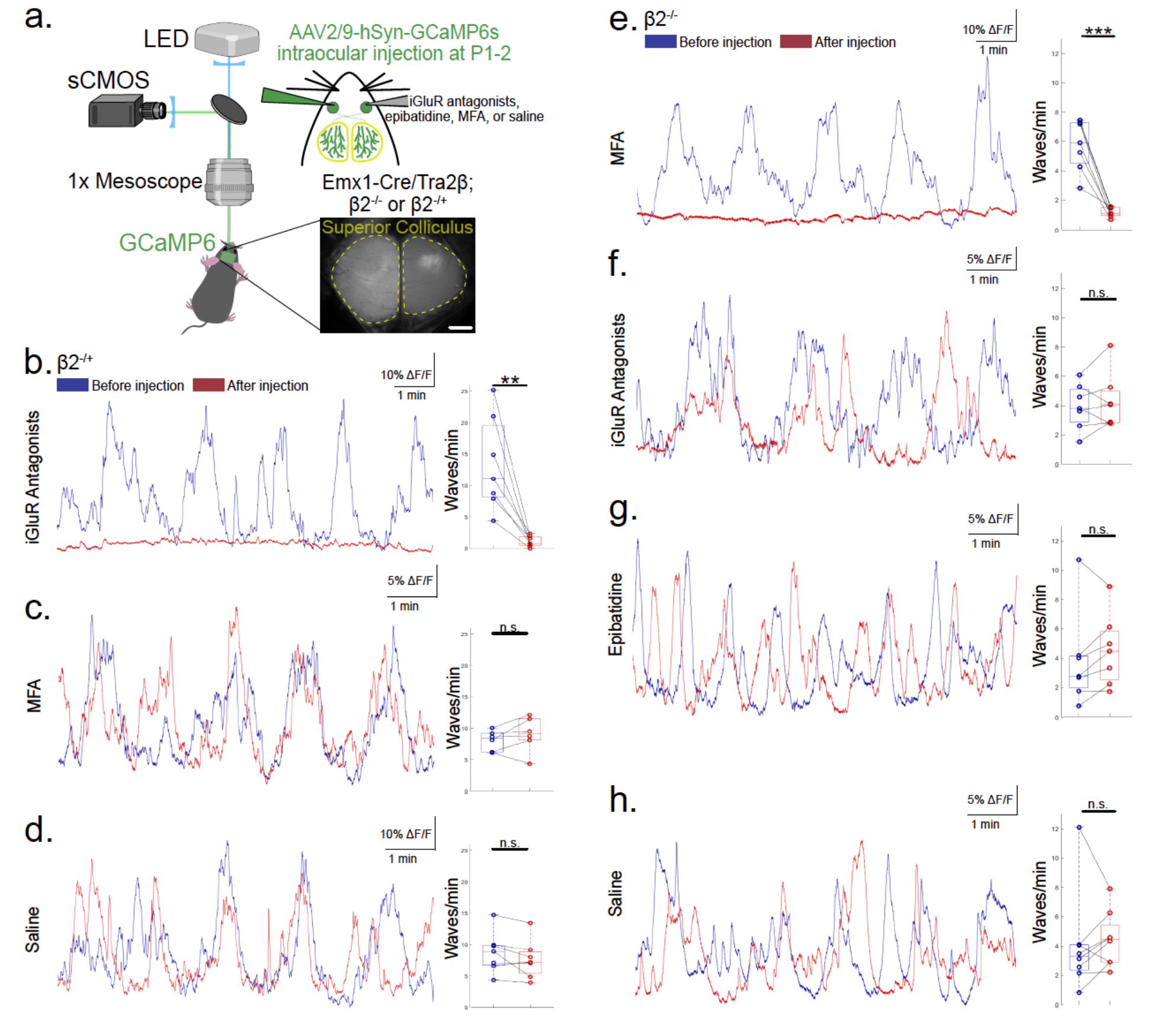
In the absence of cholinergic waves, glutamatergic waves fail to emerge *in vivo* at P9-11. **a**. Schematic of widefield single-photon microscopy used to perform calcium imaging of retinal ganglion cell axon activity in the SC of P9-11 Emx1-Cre/Tra2β^f/f^ (cortex-less mice) crossed with β2^−/+^ or β2^−/−^ mice. Spontaneous activity was recorded before and after an acute intraocular injection of either a cocktail of ionotropic glutamate receptor (iGluR) antagonists, epibatidine, MFA, or saline. Scale bar, 500 μm. **b**. Left: ι1F/F traces of calcium transients averaged across all pixels from one hemisphere of the SC before (navy) and after (maroon) an acute injection of a combination of 1 mM CP 465022 and 5 mM APV in β2^−/+^ mice. Right: Number of waves detected per minute before and after the injection. Each data point represents one animal, lines connect data from the same animal. n = 7 animals; p = 0.004. **c**. Left: ι1F/F traces of calcium transients before and after an acute injection of 20 mM MFA in β2^−/+^mice. n = 6 animals; p = 0.23. **d**. Left: ι1F/F traces of calcium transients before and after an acute injection of saline in β2^−/+^ mice. n = 7 animals; p = 0.1. **e**. Left: ι1F/F traces of calcium transients before and after an acute injection of 20 mM MFA in β2^−/−^ mice. n = 7 animals; p = 4.3e^−4^. **f**. Left: ι1F/F traces of calcium transients before and after an acute injection of a combination of 1 mM CP 465022 and 5 mM APV in β2^−/−^ mice. n = 7 animals; p = 0.44. **g**. Left: ι1F/F traces of calcium transients before and after an acute injection of 100 μM epibatidine in β2^−/−^ mice. n = 7 animals; p = 0.2. **h**. Left: ι1F/F traces of calcium transients before and after an acute injection of saline in β2^−/−^ mice. n = 8 animals; p = 0.61. All statistical tests here are paired t-tests.

### Residual gap junction-mediated activity at P9-11 *in vivo* in the absence of cholinergic waves is highly abnormal

The spatiotemporal properties of Stage III glutamatergic retinal waves have been well-documented *in vivo*. Whereas Stage II cholinergic retinal waves have large and continuous wavefronts, Stage III waves are smaller and occur more frequently than Stage II waves (Gribizis et al., 2019). Moreover, a strong temporal-to-nasal bias in the propagation direction of retinal waves emerges in late Stage II waves (P8) and persists throughout Stage III waves until shortly before eye-opening (Ge et al., 2021). To test whether the residual gap junction-mediated activity in the absence of cholinergic waves preserved any of the spatiotemporal characteristics of Stage III glutamatergic retinal waves, we compared the properties of the activity observed in β2^−/−^ mice and their littermate β2^−/+^ controls (Video 3-4 and Figure 3A).

**Fig. 3.**
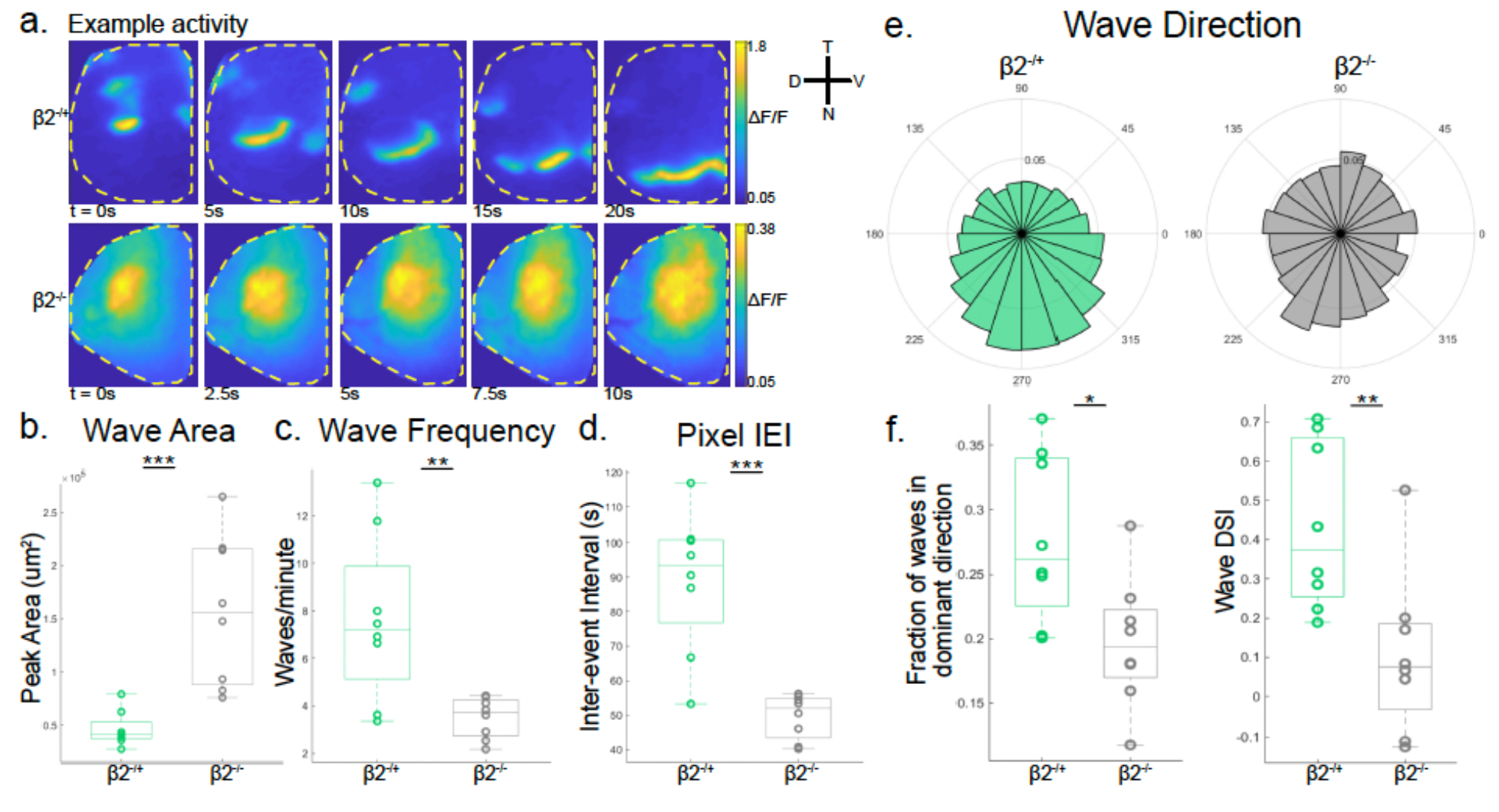
In the absence of cholinergic waves, residual P9-11 activity is highly abnormal. **a**. βF/F montages of example spontaneous events at P9-11 in β2^−/+^ (top) and β2^−/−^ (bottom) mice. **b**. Peak area of detected waves. Each data point represents one animal; n = 8 β2^−/+^ mice and n = 8 β2^−/−^ mice; p = 6.7e^−4^. **c**. Number of waves detected per minute; p = 0.006. **d**. Inter-event interval of activity detected at each pixel; p = 1.3e^−4^. **e**. Frequency distribution of propagation directions of waves. **f**. Left: Proportion of waves traveling within 60° around the dominant temporal-to-nasal direction; p = 0.015. Right: Wave DSI; p = 0.008. All statistical tests here are unpaired t-tests.

The peak areas of gap junction-mediated spontaneous events in P9-11 β2^−/−^ mice were significantly larger than in age-matched β2^−/+^ controls (Figure 3B). The frequency of spontaneous events was also significantly reduced in β2^−/−^ mice compared to β2^−/+^ controls, quantified using either the overall number of spontaneous events detected per minute across the retina or using the inter-event interval of activity detected at a given pixel location (Figure 3C-D). Finally, although retinal waves exhibited a temporal-to-nasal bias in β2^−/+^ controls, the residual activity in β2^−/−^ mice did not exhibit a significant propagation bias when quantified using either the fraction of events propagating in the temporal-to-nasal direction or Wave DSI (Figure 3E-F). These differences in spatiotemporal characteristics continued to persist in β2^−/−^ and β2^−/+^ control mice at P12-13, suggesting that Stage III retinal waves are not simply developmentally delayed in β2^−/−^ mice but rather fail to emerge at any point prior to eye-opening (Supplementary Fig 4).

Although we observed significant differences in the spatiotemporal properties of retinal axon activity between β2^−/−^ mice and β2^−/+^ controls at P9-11 *in vivo*, previous *in vitro* work examining activity patterns in these mice at the same age using two-photon calcium imaging of the retina demonstrated only a significant reduction in the frequency of events in β2^−/−^ mice but no difference in the temporal-to-nasal propagation bias in β2^−/−^ mice compared to β2^−/+^ controls (Tiriac et al., 2022). To determine whether these differences might be due to observing activity in the retina rather than the retinal axons projecting to the SC, we performed *in situ* retinal imaging of awake β2^−/−^ and β2^−/+^ mice (Wang et al, bioRxiv). When examining retinal activity directly, the differences observed in the spatiotemporal activity patterns between β2^−/−^ and β2^−/+^ mice (Stage III, P9-P11) were much less pronounced. Indeed, we found no significant difference between the peak area, duration, or frequency of spontaneous events detected in the retina in β2^−/−^ and β2^−/+^ mice. Further, the difference in propagation bias was also less pronounced with only a significant reduction in the Wave DSI of events in β2^−/−^ mice but no significant difference in the fraction of events propagating in the temporal-to-nasal direction (Supplementary Fig 5).

Direct imaging of the retina *in situ* has a smaller field of view in comparison to imaging of retinal axons in the SC. To investigate whether the differences in the temporal-to-nasal propagation bias of P9-11 spontaneous activity in β2^−/−^ and β2^−/+^ mice observed at the retinal axons or directly in the retina might be due to differences in the size and location of the imaging field of view, we restricted our analysis of spatiotemporal activity in retinal axons projecting the SC to correspond to different restricted retinal locations. *In vitro* two-photon calcium imaging of β2^−/−^ and β2^−/+^ retinas that found no difference in propagation bias was performed in the ventral retina (Tiriac et al., 2022). Thus, we restricted our analysis to the retinotopically matched area of the SC corresponding to either the dorsal or ventral retina (Seabrook et al., 2017). We observed a significant reduction in the temporal-to-nasal propagation bias of spontaneous events in β2^−/−^ mice compared to β2^−/+^ controls in the area corresponding to the dorsal, but not ventral retina (Supplementary Fig 6A-F). *In situ* retinal imaging of activity in β2^−/−^ and β2^−/+^ mice is limited to the retinal area adjacent to the optic nerve head (Wang et al, bioRxiv). To determine whether the lack of a strong propagation bias observed in retinal imaging of β2^−/+^ mice was due to the limited imaging field of view in the retina, we restricted our analysis of retinal axon activity to an area corresponding to the optic nerve head in the retina (see Methods and Materials). In the area around the optic nerve head, we found that the temporal-to-nasal propagation bias in retinal axon activity of β2^−/+^ mice was reduced, resulting in a significant difference between β2^−/−^ and β2^−/+^ mice only when using the Wave DSI but not the fraction of events propagating in the temporal-to-nasal direction (Supplementary Fig 6G-L). Taken together, the results of these analyses suggest that the ability to accurately measure the propagation bias of spontaneous retinal events depends upon the size and location of the imaging field of view, both in the retina itself and in the retinal axons projecting to the SC. With access to the full extent of the SC, and therefore the full extent of the retina in mice that develop without a cortex, significant differences in the spatiotemporal properties of spontaneous activity at P9-11 in the absence of cholinergic waves are apparent (Figure 3).

### Acute pharmacological blockade of β2-nAChR disrupts Stage III wave directionality

Our results thus far indicate that the absence of the cholinergic network in mutant β2^−/−^ mice prevents the emergence of Stage III glutamatergic retinal waves and that the compensatory activity observed at P9-11 lacks the spatiotemporal characteristics of Stage III retinal waves normally observed at the same age. We next investigate whether acetylcholine was necessary for any of the typical spatiotemporal properties characteristic of Stage III retinal waves in WT mice. To answer this question, we performed *in vivo* calcium imaging of retinal axons in P9-11 Emx1-Cre; Tra2β^f/f^ mice before and after intraocular injections of epibatidine to acutely block β2-nAChRs (Figure 4A). We observed a strong temporal-to-nasal propagation bias in baseline P9-11 spontaneous retinal waves that was significantly reduced by application of epibatidine but not saline, measured using either the fraction of waves propagating in the temporal-to-nasal direction or the Wave DSI (Video 5 and Figure 4B-E).

**Fig. 4.**
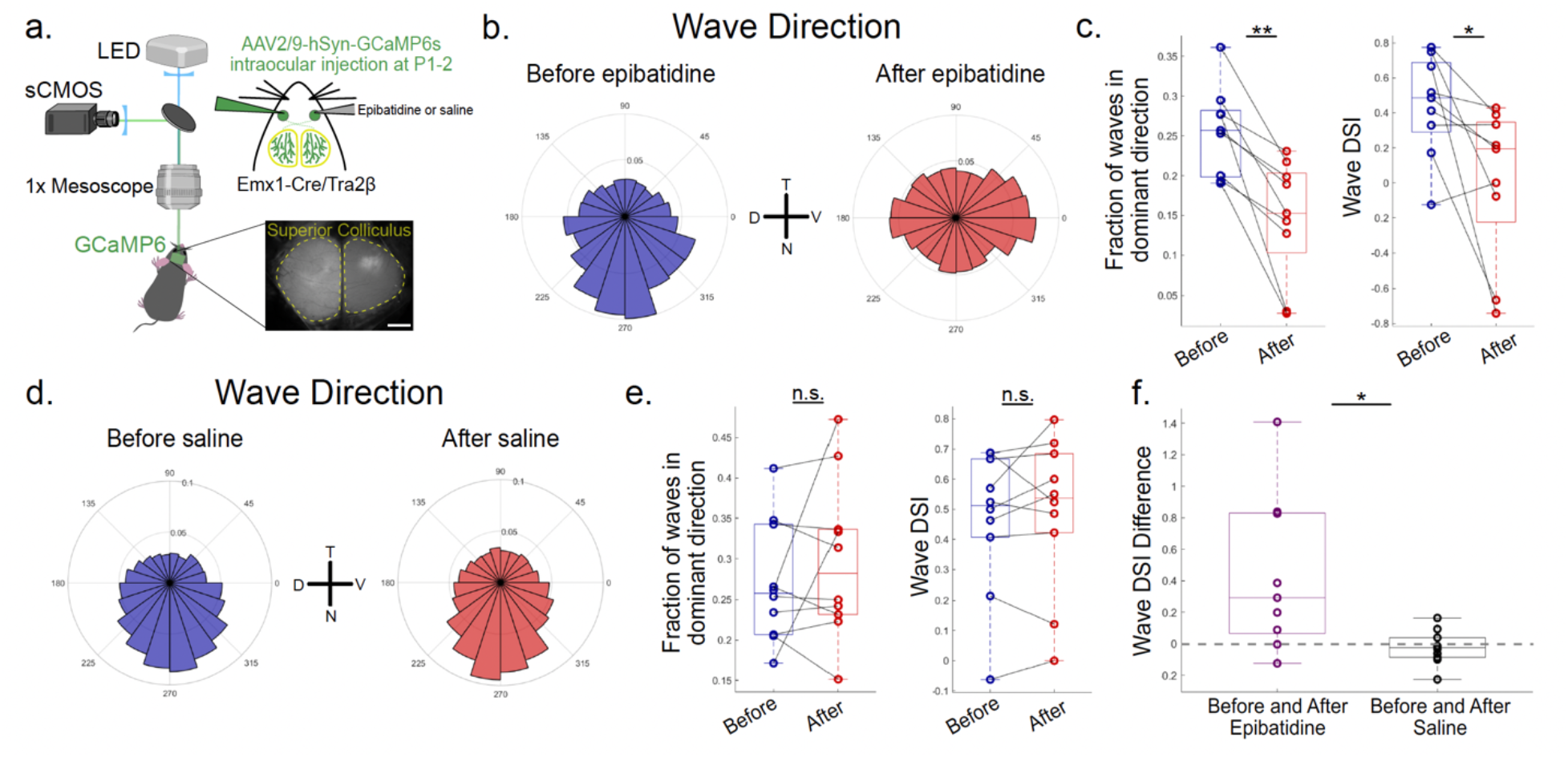
Acute pharmacological blockade of β2-nAChRs disrupts Stage III wave directionality. **a**. Schematic of widefield single-photon microscopy used to perform calcium imaging of retinal ganglion cell axon activity in the SC of P9-11 Emx1-Cre/Tra2β^f/f^ (cortex-less mice). Spontaneous activity was recorded before and after an acute intraocular injection of either epibatidine or saline. Scale bar, 500 μm. **b**. Frequency distribution of propagation directions of waves before and after an acute injection of 100 μM epibatidine; n = 9 animals. **c**. Left: Proportion of waves traveling within 60° around the dominant temporal-to-nasal direction. Each data point represents one animal; lines connect data from the same animal; p = 0.001. Right: Wave DSI; p = 0.03. Paired t-test. **d**. Frequency distribution of propagation directions of waves before and after an acute injection of saline; n = 10 animals. **e**. Left: Proportion of waves traveling within 60° around the dominant temporal-to-nasal direction; p = 0.34. Right: Wave DSI; p = 0.49. Paired t-test **f**. Difference in Wave DSI before and after an injection of 100 μM epibatidine (purple) or before and after an injection of saline (black); p = 0.01. Unpaired t-test. Dashed grey line indicates the median of a permuted null distribution calculated by shuffling the difference values 1000 times. The real median difference was significantly greater than the mean of the permuted null distribution (p = 0.025).

To further quantify the effect of epibatidine on wave directionality, we compared the change in Wave DSI before and after application of epibatidine relative to saline and found that the change in Wave DSI following saline was significantly smaller compared to the change in Wave DSI following epibatidine. We also computed a permuted null distribution by shuffling the differences in Wave DSI following either epibatidine and saline injection and found that the real observed difference in Wave DSI between epibatidine and saline was significantly greater than what could be expected by chance (Figure 4F). In addition to a significant reduction in wave directionality, we also observed that epibatidine had a significant effect on the peak wave area and distance traveled by each wave (Supplementary Fig 7).

To ensure that the effects of epibatidine on the propagation bias of Stage III retinal waves was due to the blockade of β2-nAChRs and not to potential off-target effects of epibatidine, we sought to confirm our results by utilizing DHβE, another pharmacological antagonist of β2-nAChRs. We also observed a significant reduction in the temporal-to-nasal propagation bias following an acute intraocular injection of DHβE (Supplementary Fig 8). Taken together, these results suggest that acetylcholine is a necessary circuit component underlying several spatiotemporal properties of Stage III retinal waves, including the temporal-to-nasal propagation bias. While cholinergic signaling is not necessary for the propagation of Stage III retinal waves, it plays a continued role in shaping Stage III retinal waves even after the cessation of acetylcholine-driven Stage II waves.

### Acute pharmacological blockade of α7-nAChR has no effect on Stage III wave directionality

The propagation of cholinergic retinal waves is known to specifically depend on volume transmission of acetylcholine through heteromeric β2-nAChRs (Bansal et al., 2000; Ford & Feller, 2012). However, retinal cells also express several other subtypes of ionotropic nAChRs (Ford & Feller, 2012). To further investigate the circuit mechanism underlying acetylcholine’s role in Stage III wave directionality, we next sought to test whether the pharmacological blockade of another ionotropic nAChR subtype might also influence the temporal-to-nasal propagation bias of Stage III waves. We performed intraocular injections of methyllycaconitine (MLA), an antagonist of the homomeric α7-nAChR, and measured its effect on the spatiotemporal properties of P9-11 retinal waves (Figure 5A). In contrast to the acute blockade of β2-nAChRs, MLA injection had no significant effect on Stage III wave directionality. Spontaneous retinal waves maintained a strong temporal-to-nasal propagation after both acute MLA and saline injections, suggesting that acetylcholine specifically through β2-nAChRs, but not α7-nAChRs, plays a critical role in the propagation bias of Stage III glutamatergic retinal waves (Figure 5B-E).

**Fig. 5.**
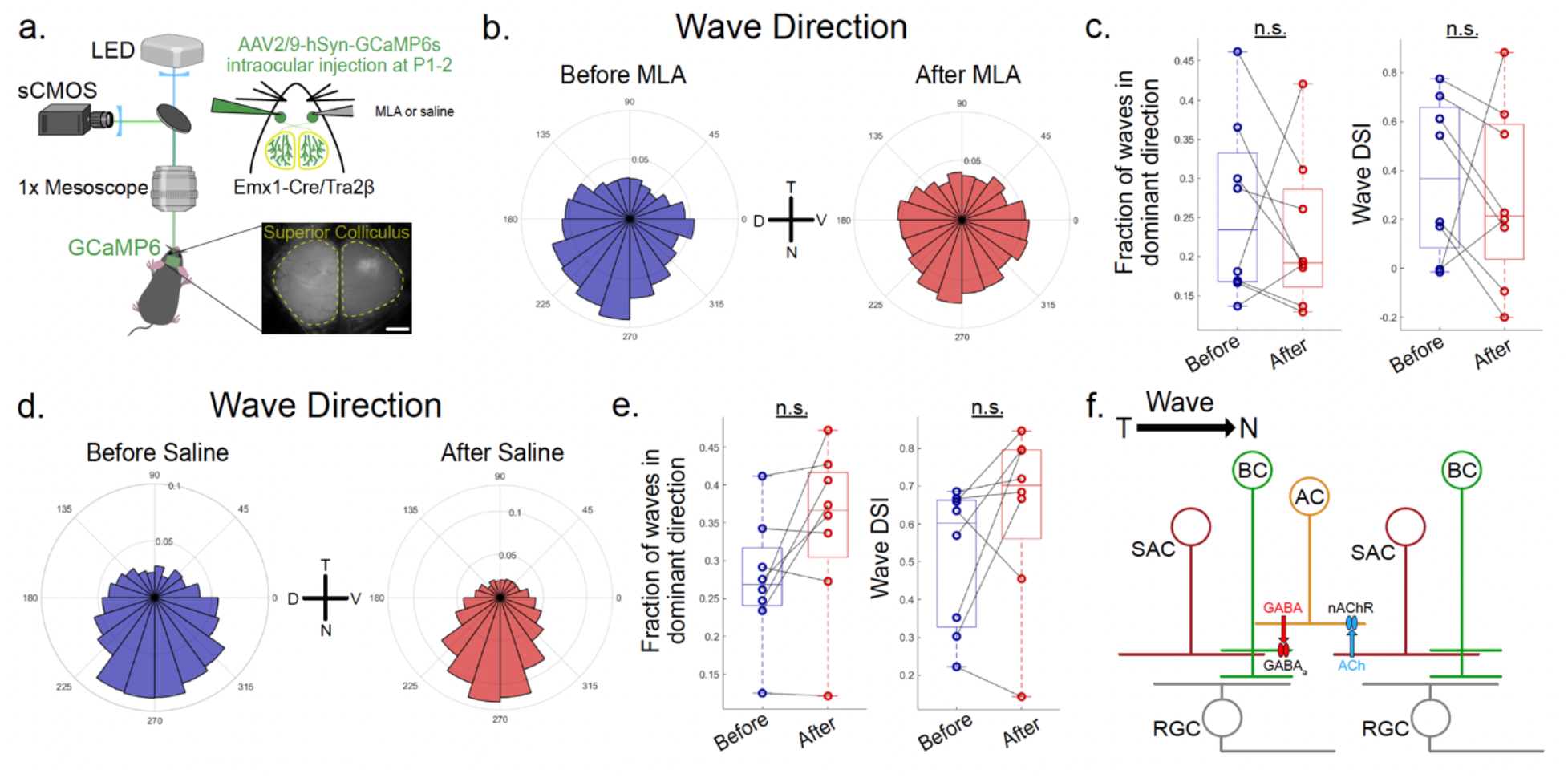
Acute pharmacological blockade of α7-nAChRs does not disrupt Stage III wave directionality. **a**. Schematic of widefield single-photon microscopy used to perform calcium imaging of retinal ganglion cell axon activity in the SC of P9-11 Emx1-Cre/Tra2β^f/f^ (cortex-less mice). Spontaneous activity was recorded before and after an acute intraocular injection of either MLA or saline. Scale bar, 500 μm. **b**. Frequency distribution of propagation directions of waves before and after an acute injection of 1μM MLA; n = 8 animals. **c**. Left: Proportion of waves traveling within 60° around the dominant temporal-to-nasal direction. Each data point represents one animal; lines connect data from the same animal; p = 0.54. Right: Wave DSI; p = 0.64. **d**. Frequency distribution of propagation directions of waves before and after an acute injection of saline; n = 8 animals.

## Discussion

We build upon previous *in vitro* work to investigate the role of the retinal cholinergic network in the transition from Stage II to Stage III retinal waves *in vivo*. We show that compensatory cholinergic activity at P9-11 in mice that lack glutamatergic waves does not display the temporal-to-nasal propagation usually observed in retinal waves at this age. Further, Stage III glutamatergic waves fail to emerge in β2^−/−^ mice that lack cholinergic waves, and in their place Stage I-like gap junction-mediated activity persists at P9-11. Moreover, the residual activity at P9-11 lacks the typical spatiotemporal properties of Stage III waves and instead occurs in large frequent bursts with no discernible propagation bias. Finally, we demonstrate that in wildtype mice, cholinergic signaling through β2-nAChRs is a critical circuit mechanism for Stage III wave directionality.

### Compensatory cholinergic activity in the absence of glutamatergic waves

Our findings replicate previously reported data showing that the cessation of each stage of retinal waves promotes the emergence of the following stage of retinal waves. *In vitro* experiments using a mutant mouse line which lacks VGLUT1 and therefore glutamatergic retinal waves found that spontaneous events recorded at P11 continued to be mediated by acetylcholine (Blankenship et al, 2009). When we assessed the pharmacological profile of spontaneous activity at P9-11 in Vsx2-SE^−/−^ mice that also lack glutamatergic waves due to the failure to develop BCs, we similarly found that the residual activity was mediated by acetylcholine *in vivo*. Taken together, these results support a model in which the emergence of the glutamatergic network facilitates the cessation of Stage II cholinergic retinal waves.

Interestingly, when we examined the spatiotemporal properties of the persisting cholinergic activity in Vsx2-SE^−/−^ mice, we found that cholinergic activity observed at P9-11 was no longer biased to propagate in the temporal-to-nasal direction. Previous findings have shown that GABAergic signaling from SACs is a required circuit element for wave directionality in Stage III retinal waves (Ge et al., 2021). Our finding further suggests that glutamatergic signaling from BCs is also required to support the temporal-to-nasal propagation bias observed in early Stage III waves. Moreover, this result suggests that persistent cholinergic activity in the absence of glutamatergic waves is not simply a continuation of Stage II retinal waves, but rather a compensatory mechanism with a unique set of spatiotemporal characteristics.

The compensatory activity preserves some features of normal Stage III retinal waves, including the size, duration, and frequency of events. Even in the absence of Stage III waves, the residual spontaneous activity may therefore provide enough correlated activity between pre– and post-synaptic targets to maintain normal visual map development. Indeed, previous *in vivo* experiments using VGLUT1 KO mice found that eye-specific segregation of retinal projections to the dLGN and SC is maintained (Xu et al., 2016). However, the lack of wave directionality in the compensatory cholinergic activity may impact downstream visual motion detection (Ge et al., 2021). Future experiments will be necessary to determine the full impact of the absence of glutamatergic waves on visual system development.

### The importance of cholinergic signaling beyond Stage II retinal waves

Both *in vivo* and *in vitro* experiments have previously demonstrated that compensatory activity mediated by gap junctions continues in the place of Stage II retinal waves in P0-7 β2^−/−^ mice (Bansal et al., 2000; Burbridge et al., 2014; Stacy et al., 2005; Sun et al., 2008). When we examined residual activity in β2^−/−^ mice at P9-11, we found that gap junction-mediated activity continues throughout the developmental window in which Stage III retinal waves would normally be observed. In contrast to previous *in vitro* work demonstrating that glutamatergic retinal waves emerge precociously in the absence of cholinergic retinal waves, we found that the compensatory activity at P9-11 *in vivo* was not blocked by iGluR antagonists, suggesting that glutamatergic waves fail to emerge entirely (Bansal et al., 2000). This discrepancy may be due to the differences between *in vivo* and *in vitro* experimental methods, but future experiments which chronically examine the developmental progress of spontaneous activity may also be helpful in elucidating whether glutamatergic activity is present at any age in β2^−/−^ mice. Our results suggest that the presence of the cholinergic network is necessary for the emergence of Stage III glutamatergic retinal waves.

In addition to its role in facilitating the emergence of Stage III retinal waves, cholinergic signaling appears to be critical for several of the typical spatiotemporal characteristics of Stage III retinal waves that are known to instruct the development of downstream retinal projections and visual neuron responses. Stage III retinal waves in ferrets require acetylcholine for the correlated retinal activity necessary to maintain eye-specific segregation in the dLGN (Davis et al., 2015). Moreover, when compensatory cholinergic activity is removed at P11 in VGLUT1 KO mice that lack Stage III waves using a conditional β2-nAChR KO in SACs, downstream eye-specific segregation in the dLGN and SC is impaired (Xu et al., 2016). Our results contribute further evidence for acetylcholine’s importance during Stage III retinal waves. When acetylcholine is disrupted either by genetic manipulation in β2^−/−^ mice or by acute pharmacological manipulation using epibatidine, we observed disruptions in the propagation bias, frequency, and size of spontaneous retinal events at P9-11 that may impair both the development of retinal projections to downstream visual structures and the development of direction-selective response in downstream SC neurons (Ge et al., 2021).

Though the role of cholinergic signaling during spontaneous retinal waves has been traditionally limited to driving Stage II retinal waves, our results demonstrate its continued importance during Stage III retinal waves. Recent work also suggests that cholinergic signaling plays an important role during Stage I retinal waves. β2-nAChR antagonism leads to a reduction in Stage I wave frequency while manipulation of other subtypes of nAChRs can block Stage I waves altogether (Voufo et al., 2023). Taken together, these results suggest that the retinal cholinergic network, including multiple subtypes of nAChRs expressed on both RGCs and SACs, plays a more complex role in mediating all stages of retinal spontaneous activity.

Intriguingly, when we observe spontaneous activity directly in the retina or indirectly in the retinal axons projecting to the SC, we find a discrepancy in the effects of acetylcholine manipulation on the spatiotemporal properties of Stage III retinal waves. Though RGC axon activity in β2^−/−^ mice was highly abnormal in its frequency, size, and propagation bias, *in situ* retinal imaging of β2^−/−^ mice did not exhibit the same differences in spatiotemporal characteristics. These discrepancies may be due to several factors. First, the imaging field of view in the retina is limited to the area directly adjacent to the optic nerve head whereas our use of mice that develop without a cortex allows access to nearly the full extent of the retina when imaging axons that project to the SC. Indeed, when we limit our analysis to a region of the SC that corresponds to the area of the retina believed to be captured by our *in situ* retinal imaging technique, we find that the difference in propagation bias between β2^−/−^ and β2^−/+^ mice is less pronounced.

Second, previous work has demonstrated that the development of retinal projections to the SC in β2^−/−^ mice is severely disrupted. Rather than refining to their retinotopic target location in the SC, retinal axon arbors in β2^−/−^ mice remain enlarged compared to WT mice at the end of the first postnatal week. The enlarged wave area we observe at P9-11 in β2^−/−^ mice may therefore reflect the abnormal development of retinocollicular projections in these mice (Dhande et al., 2011). *In situ* retinal imaging of these same mice reflects the activity of all retinal cell bodies rather than the RGC axons. Thus, while retinal cell bodies may fire in more spatially limited correlated bursts in β2^−/−^ mice, enlarged retinal axon projections to the SC might have a stronger impact on activity in downstream visual structures.

There are conflicting reports about whether direction selectivity in the retina is impaired by the β2^−/−^ mutation. While multielectrode array recordings in the retina found that the development of DSGCs was unaffected in β2^−/−^ mice, two-photon imaging of the retina found that horizontal direction selectivity was absent in β2^−/−^ mice (Elstrott et al., 2008; Tiriac et al., 2022). On the other hand, downstream visual motion detection has been shown to be impaired in β2^−/−^ mice. Specifically, both the optokinetic reflex and SC neuron direction-selective responses along the horizontal axis are severely disrupted in β2^−/−^ mice (Wang et al., 2009). Whether the downstream effects of the β2^−/−^ mutation are due to disruptions in Stage III wave directionality is unclear. Similar to our *in situ* retina recordings, two-photon imaging of the retina in β2^−/−^ mice revealed a smaller effect of the mutation on the propagation bias of Stage III retinal waves, suggesting that the wave propagation bias does not instruct the development of direction selectivity in the retina (Tiriac et al., 2022). However, the striking reduction of wave directionality measured in the retinal axons of β2^−/−^ mice along with previous evidence of disrupted SC direction selectivity in β2^−/−^ mice or in WT mice with chronically disrupted wave directionality during development suggests that the effects of the β2^−/−^ mutation could be on downstream direction-selective circuits rather than in the retina itself (Wang et al., 2009; Ge et al., 2021).

### A circuit model for Stage III wave directionality

Our results reveal an important role for acetylcholine in mediating the temporal-to-nasal propagation bias of Stage III retinal waves. Previous work has shown that the circuit mechanism for Stage III wave directionality requires asymmetric GABAergic inhibition from SACs, perhaps through feedback inhibition onto BCs. However, this model did not consider a role for acetylcholine (Ge et al., 2021). Thus, we offer an updated circuit model for Stage III wave directionality that includes cholinergic modulation (Figure 5F).

Our pharmacological manipulations suggest that acetylcholine specifically through β2– nAChRs but not α7-nAChRs is required for Stage III wave directionality. Recent work has revealed a role for acetylcholine in the modulation of the rod bipolar cell (RBC) pathway in rat retina. The RBC pathway is a well-studied circuit underlying scotopic vision in rodents in which glutamate release from RBCs onto AII amacrine cells (ACs) is controlled by reciprocal GABAergic synapses between A17 ACs and RBCs. GABAergic release from A17 ACs onto RBCs is further modulated by cholinergic signaling that is sensitive to DHβE but not MLA (Elgueta et al., 2015). These findings suggest that cholinergic transmission from SACs can influence the GABAergic inhibition of BCs through β2-nAChRs. The identification of a GABAergic A17 AC terminating onto RBCs in mouse retina suggests the existence of a similar circuit motif in the mouse (Siegert et al., 2009).

We propose a model for Stage III wave directionality in which SACs modulate the GABAergic inhibition of BCs through an intermediary AC. An asymmetry in such a circuit either at the cholinergic synapse between SACs and the intermediary AC or at the GABAergic synapse between the intermediary AC and BCs could facilitate the propagation of Stage III retinal waves in the temporal-to-nasal, but not nasal-to-temporal direction. Indeed, asymmetric crossover inhibition from ON BCs to OFF BCs through an intermediary inhibitory AC, presumed to be the AII AC, has been previously reported in Stage III retinal waves, facilitating the temporally offset sequential firing of ON followed by OFF RGCs (Akrouh and Kerchensteiner, 2013). The specific identity of the intermediary AC remains unknown.

Candidates include the A17 AC implicated in the cholinergic modulation of the RBC pathway, the AII AC implicated in crossover inhibition during Stage III retinal waves, or another yet unidentified AC that receives cholinergic input from SACs and provides GABAergic inhibition onto BCs. For example, a circuit motif involving SAC to widefield AC to BC connectivity was recently identified as playing a role in the contextual modulation of retinal direction selectivity (Huang et al., 2019). As the function of many widefield ACs remains unknown, future work is required to further substantiate our proposed model for acetylcholine’s role in mediating Stage III wave directionality.

Taken together, the results of our study show the importance of acetylcholine in spontaneous retinal activity far beyond Stage II retinal waves. Indeed, the cholinergic network continues to play a vital role both in the emergence of Stage III retinal waves and the spatiotemporal properties of Stage III retinal waves that instruct downstream visual system development. Future investigation of retinal cholinergic circuits during development will inform a deeper understanding of the evolution of acetylcholine’s role throughout all stages of retinal waves.

## Materials and Methods

### Animals

Animal care and use followed the Yale Institutional Animal Care and Use Committee (IACUC), the US Department of Health and Human Services, and institution guidelines. All studies employed a mixture of male and female mice. Emx1-Cre (stock number: 027784) were purchased from the Allen Institute. Tra2β^f/f^ mice were provided by D.A. Feldheim and have been described previously (Ge et al., 2021; Shanks et al., 2016). Vsx2 CRC-SE KO mice were provided by M. A. Dyer and have been described previously (Gamlin et al., 2020; Norrie et al., 2019). Homozygous and heterozygous and wildtype littermate controls were used. We used a line of floxed β2-nAChR mice previously generated by our lab to specifically delete β2-nAChR2. β2 KO mice were generated and genotyped according to previously published strategies (Burbridge et al., 2014; Xu et al., 2009). Homozygous and heterozygous littermate controls were used. All mutant mouse lines were crossed with cortex-less Emx1-Cre/Tra2β^f/f^ mice to gain access to the full extent of the SC.

### Eye Injections

P1-P2 whole eye fill was performed as previously described (Ackman et al., 2012; Ge et al., 2021). Mouse pups were anesthetized using ice. 500 nL of AAV2/9-hSyn-GCaMP6s (Addgene, titer ≥ 1×10^13^ vg/mL) was pressure injected into each ocular vitreous through a glass micropipette using a Nanoject (Drumond Scientific). After injection, mouse pups were allowed to recover from anesthesia on a heating pad and were then returned to their mothers.

For acute pharmacological manipulation *in vivo* at P9-11, animals were re-anesthetized after 60 minutes of imaging. Eyes were exposed after 5-10 min of 2% isoflurane. 1 uL of either a cocktail of 1mM CP 465022 (Tocris) and 5 mM AP5 (Tocris), 100 uM epibatidine (Sigma-Aldrich), 20 mM MFA (Sigma-Aldrich), 1 uM MLA (Tocris), 200 uM DHβE (Tocris), or saline was pressure injected into the vitreous. After injection, the eyelid was closed and covered in ophthalmic ointment and a brief local anesthetic. Animals were allowed to recover from anesthesia for 20-30 minutes before being returned to imaging.

### Surgical procedure for in vivo imaging of the SC

All surgeries were performed in accordance with the regulations set by the Yale University IACUC and in accordance with NIH guidelines. Mice were surgically prepared for imaging as previously reported (Ackman et al., 2009; Ge et al., 2021). Briefly, mice were anesthetized with isoflurane (2-2.5%) via a nose cone and placed on a heating pad set to 37°C (HTP-1500, Adroit). Carprofen was administered subcutaneously (5 mg/kg). Local anesthesia was provided by subcutaneous injection (0.02-0.05 mL) of 1% xylocaine (10mg/mL lidocaine per 0.01 mg/mL epinephrine, AstraZeneca) under the scalp. After removal of the scalp, steel screws with flat heads were fixed to the exposed skull using a thin layer of Vetbond (3M) followed by dental cement (Metabond, Parkell). Isoflurane anesthesia was adjusted between 0.5-1.0% as necessary to maintain a stable respiratory rate. An approximately 4mm oval craniotomy was performed above the superior colliculus and the dura was carefully removed. A double-layer glass coverslip was placed and secured over the surface of the tissue using cyanoacrylate glue. After surgery, mice were allowed to recover for 1.5-4 hours on a heating pad.

### Surgical procedure for in situ retinal imaging

All surgeries were performed in accordance with the regulations set by the Yale University IACUC and in accordance with NIH guidelines. Mice were surgically prepared for imaging as previously reported (Wang et al., bioRxiv). Briefly, mice were anesthetized with isoflurane (2-3%) via a nose cone and placed on a heating pad set to 37°C (HTP-1500, Adroit). Carprofen was administered subcutaneously (5 mg/kg). Local anesthesia was provided topically using 0.5% lidocaine. Approximately 2mm of the scalp was removed to expose the skull and two steel head posts were attached to the skull with cyanoacrylate glue. The eyelid and approximately 0.5mm of the surrounding skin were carefully removed from the eye. Drops of atropine sulfate (1%, Millipore Sigma) were administered to dilate the pupil followed by drops of hypromellose (2.5% dissolved in 1× PBS, Millipore Sigma) to maintain a moist environment. The extraocular muscles were removed and approximately 10 uL of Vetbond (3M) was injected behind the eyeball to fix the eyeball to the orbit. The eyeball was further fixed from above with a layer of cyanoacrylate glue, covering the exposed outer area of the eyeball and surrounding skull, ensuring that the pupil remained clear and unobstructed. The exposed pupil was then covered with hypromellose followed by a round 5mm glass coverslip. The coverslip was secured using cyanoacrylate glue. After surgery, mice were allowed to recover on a heating pad with an oxygen supply for 1.5 hours before imaging.

### Widefield single-photon calcium imaging

Widefield calcium imaging of both the SC and the retina was performed using a Zeiss AxioZoom v.16 microscope with a PlanNeoFluar Z 1×/0.25 objective. Illumination was provided by an LED source (X-Cite CLED1) with blue light (470 nm, Chroma ET470/20×) for GCaMP imaging. Fluorescence emissions were passed through a dichroic (Chroma T4951pxr) and an emission filter (Chroma ET525/50m) and was collected by a sCMOS camera (pco.edge 4.2, PCO) affixed to the microscope. Images were collected by Camware software at 10 Hz.

Imaging was performed on unanesthetized, head-fixed mice in a dark box after 1.5-4 hours recovery from the surgical procedure. For retinal imaging, the retina was allowed 10min under consistent blue light illumination for adaptation. Each recording consisted of a single, continuously acquired movie for 10 min. At least 6 recordings per animal were collected, and at least 4 recordings per animal following an acute pharmacological manipulation were collected.

### Widefield calcium imaging preprocessing and wave detection

Raw TIFF movies were pre-processed using methods adapted from Wang et al., biorxiv and Ge et al., 2021. Each movie was downsampled (average-pooling with a 2×2 filter) and corrected for motion using subpixel rigid registration (Pnevmatikakis & Giovannuci, 2017). ROI masks were manually defined in ImageJ prior to pre-processing. Photobleaching correction was performed using a single-term exponential fit conducted on mean fluorescent traces of tiling 4×4 squares. Gaussian smoothing with a sigma/filter size of size = 1 was performed, followed by denoising based on singular-vector decomposition preserving the top 256 dimensions. Finally, fluorescence values were normalized by calculating the ι1F/F where F0 was defined as the 5^th^ percentile value of each pixel. Key parameters were kept consistent across conditions.

### Measurement of wave spatiotemporal properties

Automated image segmentation and calcium event detection was performed using custom routines written in MATLAB. Pre-processed ι1F/F movies were z-scored and then binarized by manually selecting a threshold for n std + mean (ranging from threshold = 2 to threshold = 5 based on signal to noise quality of the raw TIFF) for each pixel to generate a binarized movie. Individual waves (active 3D calcium domains) were automatically segmented as contiguously connected components in space and time from this binarized movie. A set of regional measurements, such as duration, diameter, centroid, and area, were obtained from function ‘regionprops’ in the MATLAB Image Processing Toolbox. Domains having a duration of less than 2 frames or an area of less than 5 pixels were ignored. Key parameters were kept consistent across conditions.

Pixel activity intervals were determined as the set of time intervals between the offset of previous activity and the onset of sequential activity for each pixel. Wave durations were determined by the number of contiguous frames for each segmented 3D domain. The largest area of a single frame for each 3D domain was taken as the peak area for each wave. Wave propagation distances were quantified as the cumulative distance between centroids of sequential frames for each wave. Wave frequencies were determined by the number of waves divided by the length of time for each recording. Wave directions were determined by calculating the vector sum of optic flow between sequential frames for individual pixels in each wave, described in greater detail in Ge et al., 2021. In brief, the propagation directions of valid waves were extracted using the “estimateFlow” MATLAB function based on the Lucas-Kanade method (Baker & Matthews, 2004). The optic flow experienced by all active pixels at each time point was pooled to determine the wave motion vector between sequential frames. A single wave direction was assigned by averaging the wave motion vector within the active duration of the wave. Wave DSI was calculated as the difference between the proportion of waves traveling within 60° around the dominant temporal-to-nasal direction and the proportion of waves traveling within 60° around the opposite nasal-to-temporal direction, divided by the sum.

To restrict the ROI for analysis in Figure 3 – figure supplement 3, an additional mask was superimposed on each SC hemisphere to remove pixels outside the ROI for wave detection and further analysis. We referred to Seabrook et al., 2017 to determine the area of the SC that was retinotopically matched to the dorsal or ventral retina, respectively. To restrict the analysis area of the SC to the area adjacent to the optic nerve in the retina, we used two different methods. For “optic disc method 1”, we estimated the retinotopically matched location in the SC using previously reported results of retrograde fluorescence tracing of retinal projections to the SC (Ellis et al., 2016). When retrograde DiI and DiD injections were performed in the medial and lateral edges of the SC, the optic nerve head was located approximately in the middle between the labeled RGCs on opposite sides of the retina. When retrograde injections of fluorescence tracers were performed in the rostral and caudal edges of the SC, the labeled RGCs projecting to the rostral SC were much closer to the optic nerve head than RGCs projecting to the caudal SC. We thus reasoned that the location corresponding to the optic nerve head in the SC would lie at the center of the medial-lateral axis and biased toward the rostral end of the rostral-caudal axis in the SC. To determine a size of the restricted field of view in the SC that would be proportional to the size of the imaging field of view in the retina, we first estimated that the average mouse retinal surface was approximately 17 mm^2^ based on reports in the literature ranging from 15-19 mm^2^. We measured the average size of the retinal field of view during *in situ* retinal imaging to be approximately 0.79 mm^2^, representing approximately 4.6% of the total retinal area. We measured that the average surface area of one hemisphere of the SC was approximately 3.13 mm^2^. A field of view corresponding to 4.6% of the total SC area is therefore approximately 0.14 mm^2^. We thus drew a mask corresponding to this size. For “optic disc method 2”, we observed that in all our animals, there was a bright patch of fluorescence expression approximately in the center of each hemisphere of the SC. We reasoned that our method of intraocular injection of viral expression GCaMP6s might result in an aggregation of fluorescence at the optic nerve head where RGC axons converge and exit the retina, thus similarly resulting in a patch of increased fluorescence expression at the location where RGC axons invade the SC. We thus drew a mask around the area of increased fluorescence expression in each hemisphere of the SC.

### Statistical analysis

Data were analyzed and plotted using custom routines implemented in MATLAB. Data from the same animal before and after an acute pharmacological manipulation were compared using paired Student’s t-tests. Data from different groups of mice were compared using unpaired Student’s t-tests. For the permutation test reported in Figure 4, we also computed a permuted null distribution by shuffling the differences in Wave DSI following either epibatidine and saline injection. Difference values were randomly assigned either to epibatidine or saline injections, and shuffling was performed 1000 times to compute the null distribution. All reported “n” refer to the number of animals and all boxplots report medians with the 75% confidence interval. Stars in figures indicate the following: * = p < 0.05, ** = p < 0.01, *** = p < 0.001.

## Acknowledgments

We thank D. Clark, J. Demb, M. Higley, L. Liang, and all members of the Crair lab for their helpful comments on this project; X. Ge and Y. Wang for help with data analysis code; M. A. Dyer for providing mouse lines; and especially Y. Zhang for help with mouse husbandry and genotyping.

## Author contributions

K.Z. and M.C.C. conceived and designed the study. K.Z. performed and analyzed wide-field calcium imaging experiments in the SC. A.S. and Y.W. performed wide-field calcium imaging experiments in the retina. K.Z. and M.C.C. analyzed and interpreted the results. K.Z. wrote the manuscript with editorial input from M.C.C. and feedback from all authors.

## Data and materials availability

All data are available in the main text or the supplementary materials.

---

**Supplementary Fig. 1.**
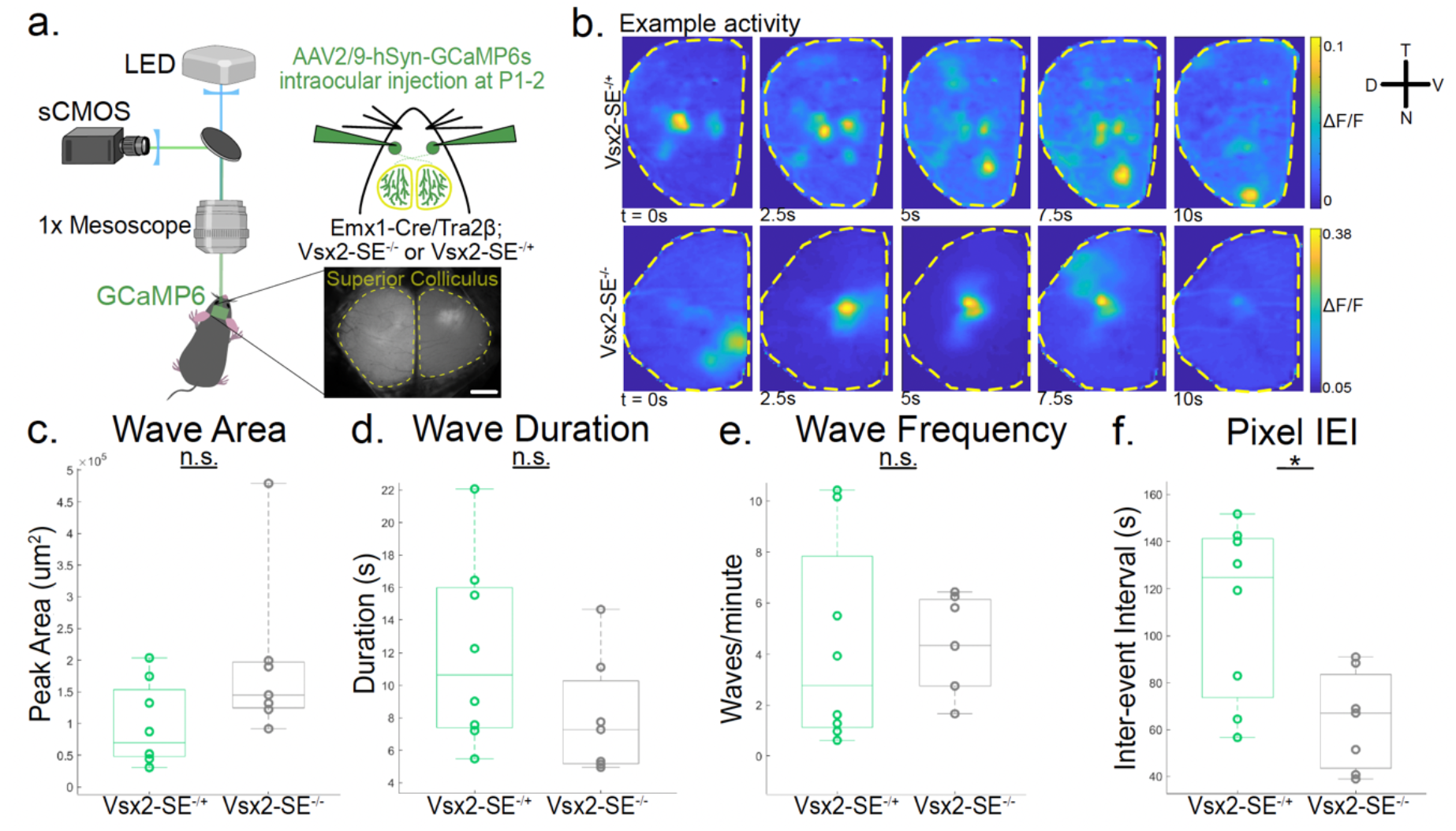
Other spatiotemporal properties of residual P9-11 activity in the absence of glutamatergic waves (related to Fig. 1). **a**. Schematic of widefield single-photon microscopy used to perform calcium imaging of retinal ganglion cell axon activity in the SC. GCaMP6s expression in retinal ganglion cells was achieved by intraocular injections of AAV2/9-hSyn-GCaMP6 at P1 or P2 in Emx1-Cre/Tra2β^f/f^ (cortex-less mice) crossed with Vsx2-SE^−/−^ or Vsx2-SE^−/+^ mice. Scale bar, 500 μm. **b**. βF/F montages of example waves at P9-11 in Vsx2-SE^−/+^ (top) and Vsx2-SE^−/−^ (bottom) mice. **c**. Peak area of detected waves. Each data point represents one animal; n = 8 Vsx2-SE^−/+^ mice and n = 7 Vsx2-SE^−/−^ mice; p = 0.09. **d**. Median duration of detected waves; p = 0.14. **e**. Number of waves detected per minute; p = 0.98. **f**. Inter-event interval of activity detected at each pixel; p = 0.01. All statistical tests here are unpaired t-tests.

**Supplementary Fig. 2.**
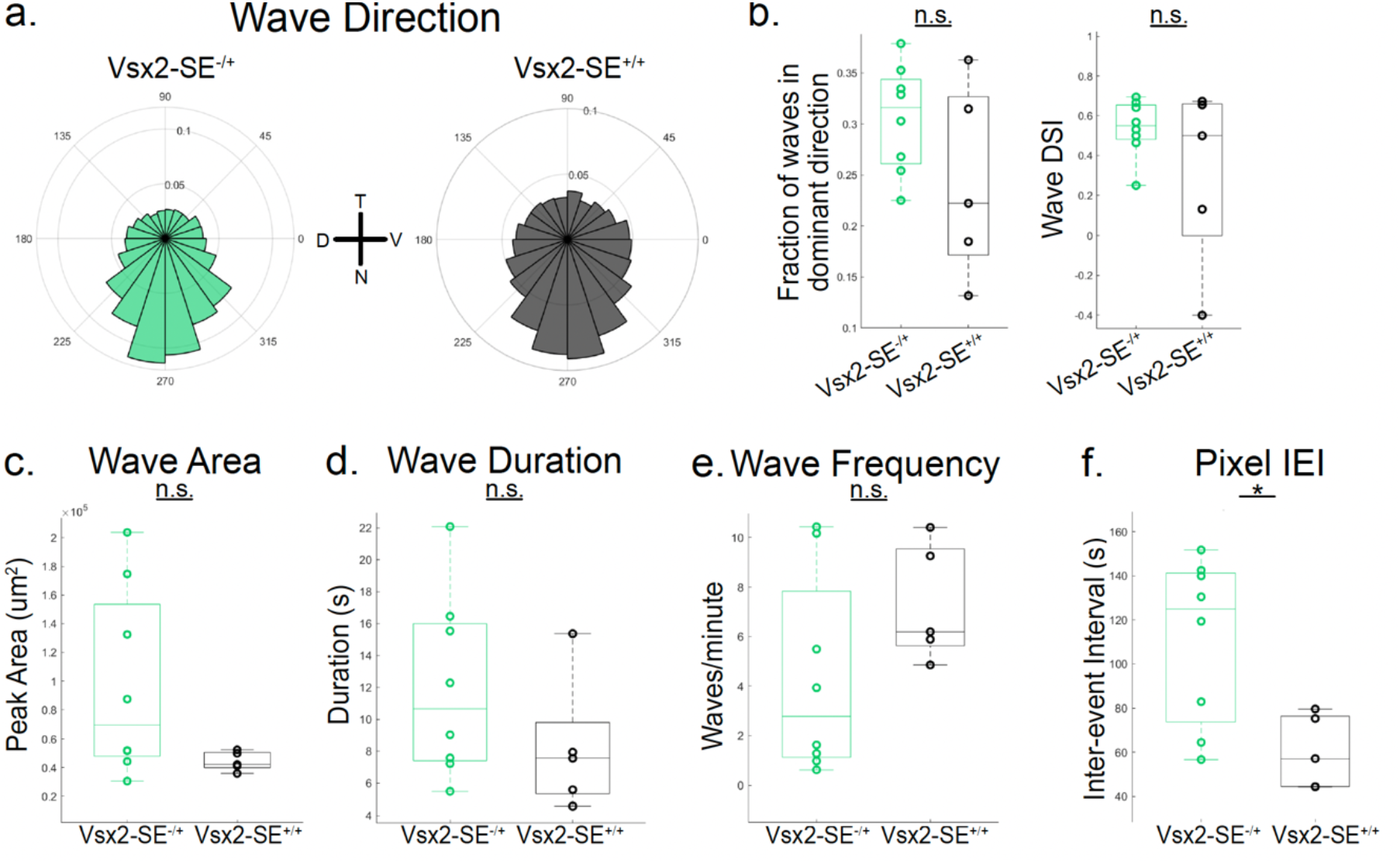
Comparison of Vsx2-SE^−/+^ and Vsx2-SE^+/+^ activity patterns at P9-11 (related to Fig. 1). **a**. Frequency distribution of propagation directions of waves in P9-11 Vsx2-SE^−/+^ (n = 8 animals; same data as in Figure 1 and Supplementary Fig 1) and Vsx2-SE^+/+^ mice (n = 5 animals). **b**. Left: Proportion of waves traveling within 60° around the dominant temporal-to-nasal direction. Each data point represents one animal; p = 0.15. Right: Wave DSI; p = 0.21. **c**. Peak area of detected waves; p = 0.11. **d**. Median duration of detected waves; p = 0.23. **e**. Number of waves detected per minute; p = 0.16. **f**. Inter-event interval of activity detected at each pixel; p = 0.02. All statistical tests here are unpaired t-tests.

**Supplementary Fig. 3.**
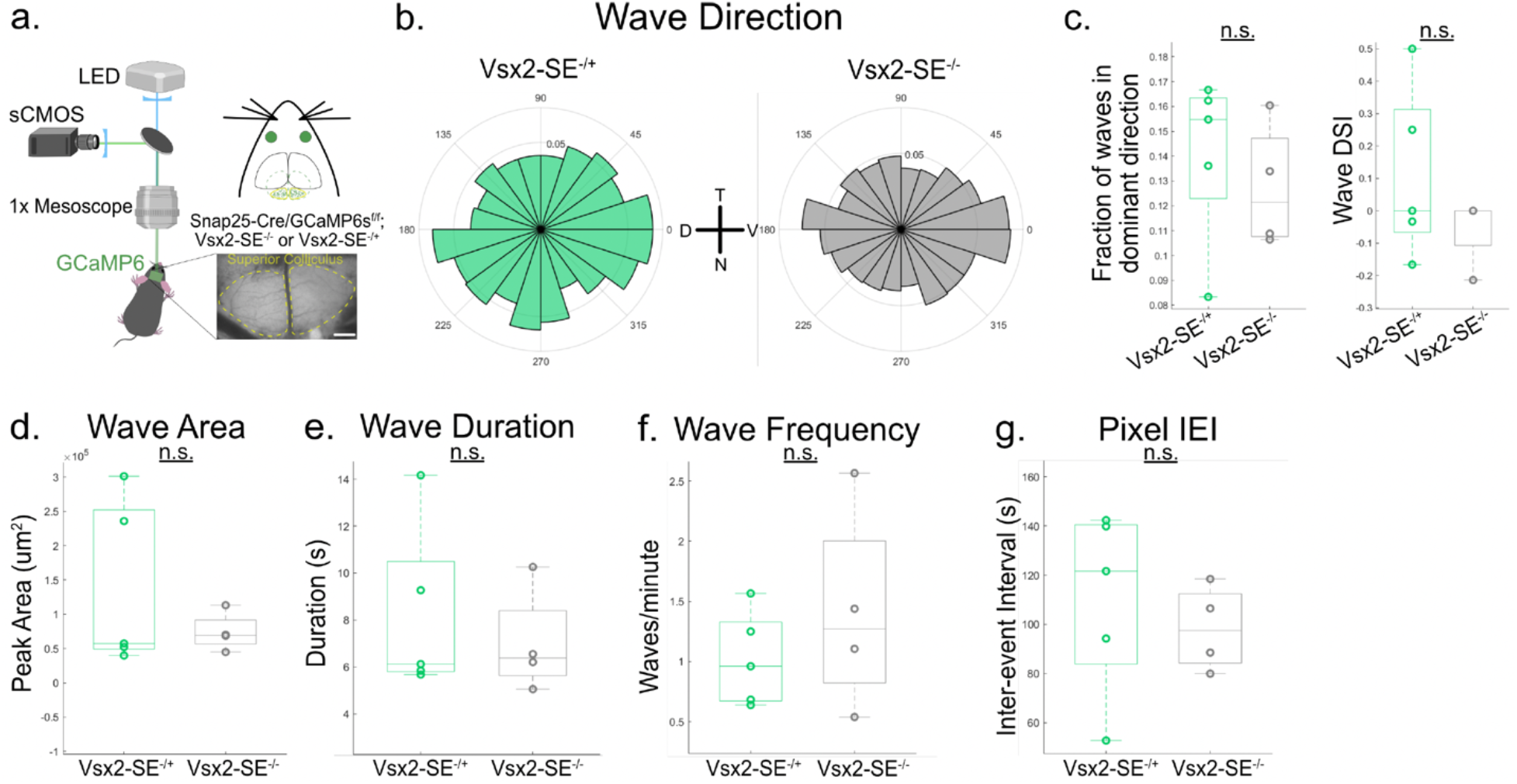
Stage II retinal wave characteristics in P5-7 Vsx2-SE^−/+^ and Vsx2-SE^−/−^ mice (related to Fig. 1). **a**. Schematic of widefield single-photon microscopy used to perform calcium imaging of retinal ganglion cell axon activity in the SC of P5-7 Snap25-Cre/GCaMP6s^f/f^ crossed with Vsx2-SE^−/+^ or Vsx2-SE^−/−^ mice. **b**. Frequency distribution of propagation directions of waves in P5-7 Vsx2-SE^−/+^ (n = 5 animals) and Vsx2-SE^−/−^ mice (n = 4 animals). **c**. Left: Proportion of waves traveling within 60° around the dominant temporal-to-nasal direction. Each data point represents one animal; p = 0.54. Right: Wave DSI; p = 0.29. **d**. Peak area of detected waves; p = 0.35. **e**. Median duration of detected waves; p = 0.59. **f**. Number of waves detected per minute; p = 0.39. **g**. Inter-event interval of activity detected at each pixel; p = 0.58. All statistical tests here are unpaired t-tests.

**Supplementary Fig. 4.**
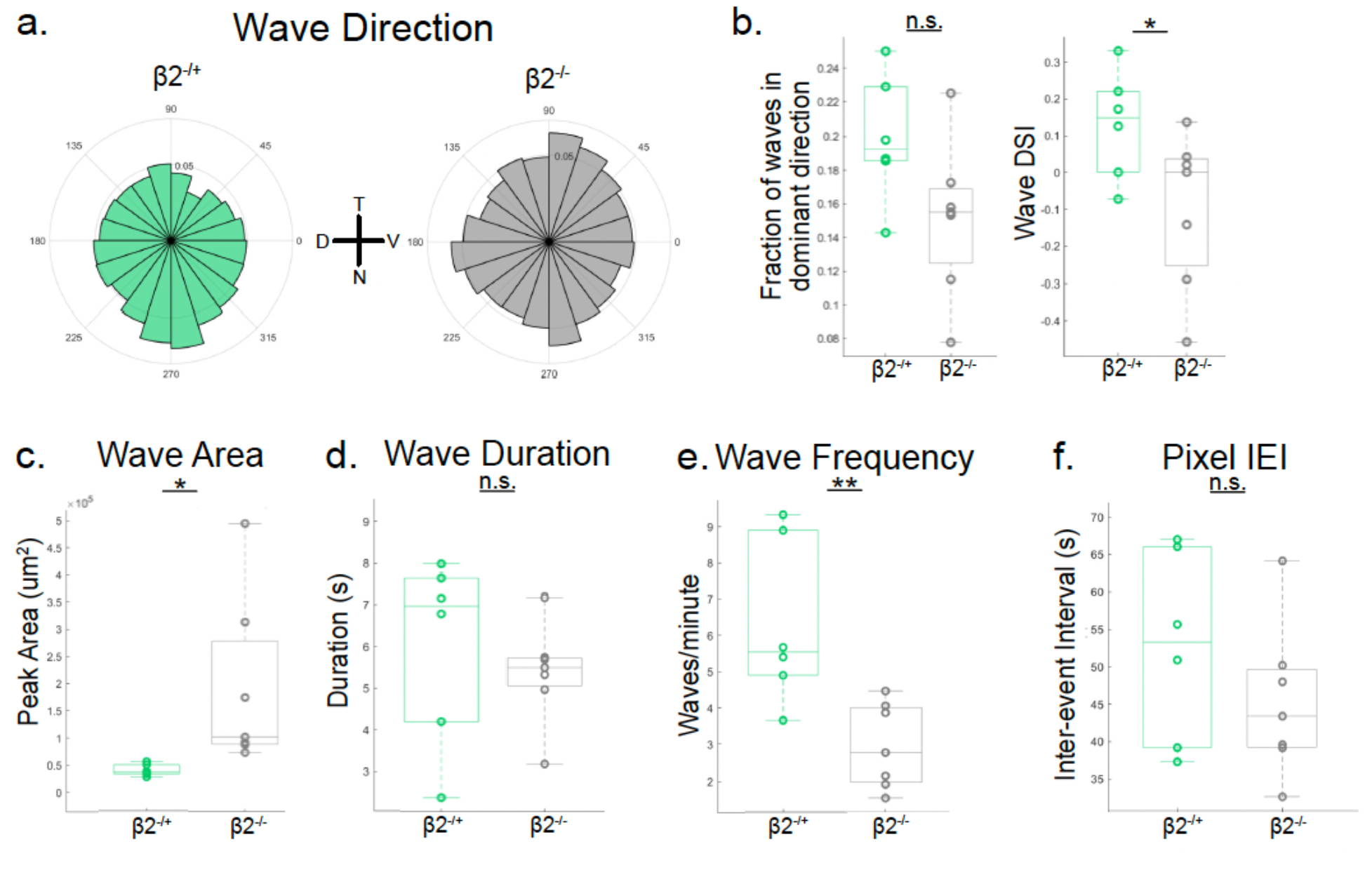
Residual activity in β2^−/−^ mice at P12-13 (related to Fig. 3). **a**. Frequency distribution of propagation directions of waves in P12-13 β2^−/+^ (n = 6 animals) and β2^−/−^ mice (n = 7 animals). **b**. Left: Proportion of waves traveling within 60° around the dominant temporal-to-nasal direction; p = 0.07. Right: Wave DSI; p = 0.048. **c**. Peak area of detected waves; p = 0.04. **d**. Median duration of detected waves; p = 0.52. **e**. Number of waves detected per minute; p = 0.006. **f**. Inter-event interval of activity detected at each pixel; p = 0.27. All statistical tests here are unpaired t-tests.

**Supplementary Fig. 5.**
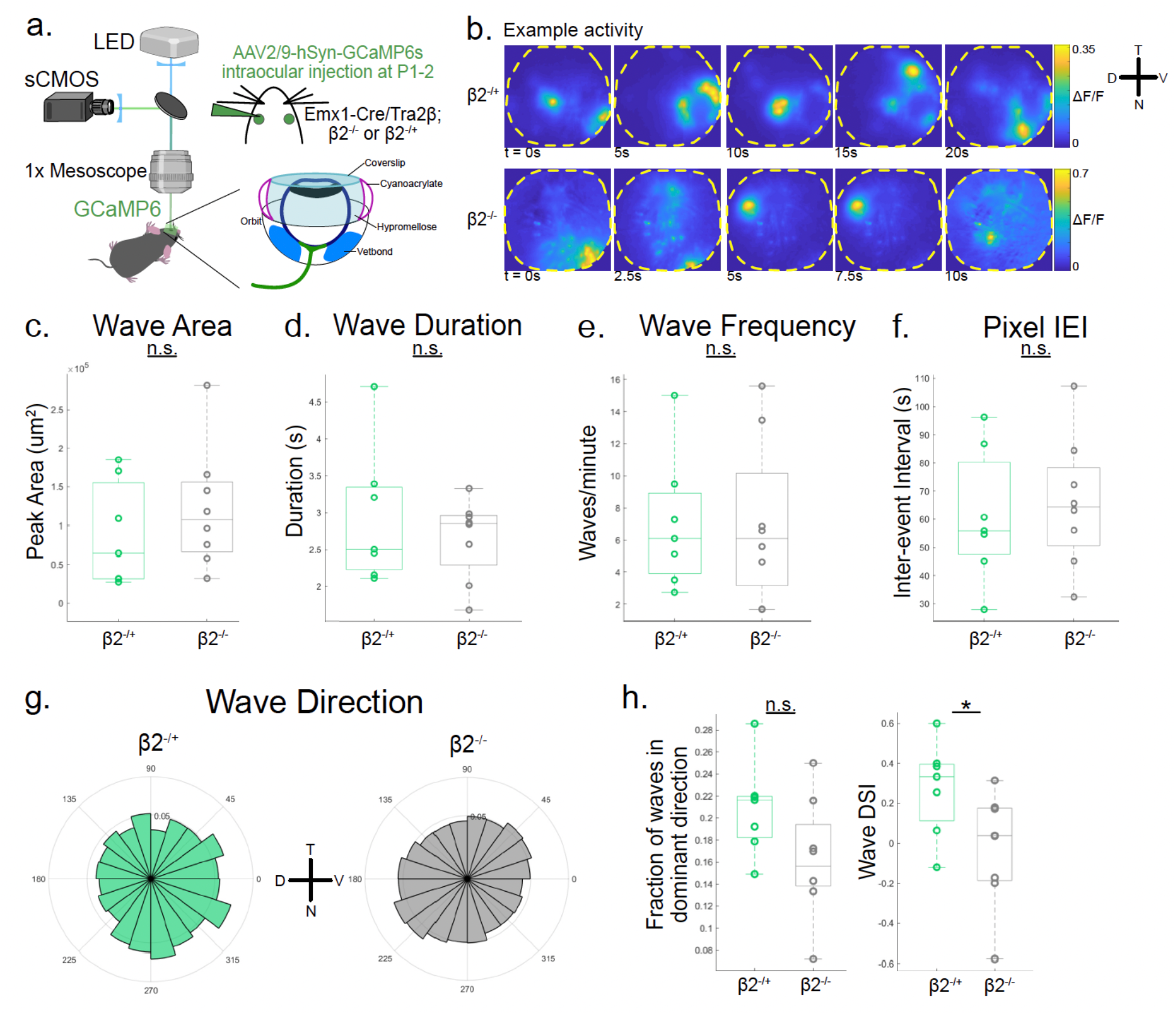
*In situ* retinal imaging of β2^−/+^ and β2^−/−^ mice at P9-11 (related to Fig. 3). **a**. Schematic of widefield single-photon microscopy used to perform calcium imaging of retinal activity through the pupil. GCaMP6s expression in retinal cells was achieved by intraocular injections of AAV2/9-hSyn-GCaMP6 at P1 or P2 in Emx1-Cre/Tra2β^f/f^ (cortex-less mice) crossed with β2^−/+^ or β2^−/−^ mice. An optical window was surgically prepared before imaging at P9-11 (see Methods for details). Mice were unanesthetized and head-fixed during imaging. Spontaneous activity was recorded for 90 minutes in each animal. **b**. βF/F montages of example retinal waves imaged directly in the retina at P9-11 in β2^−/+^ (top) and β2^−/−^ (bottom) mice. **c**. Peak area of detected waves in β2^−/+^ (n = 7 animals) and β2^−/−^ mice (n = 8 animals). Each data point represents one animal; p = 0.4. **d**. Median duration of detected waves; p = 0.48. **e**. Number of waves detected per minute; p = 0.99. **f**. Inter-event interval of activity detected at each pixel; p = 0.7. **g**. Frequency distribution of propagation directions of waves. **h**. Left: Proportion of waves traveling within 60° around the dominant temporal-to-nasal direction; p = 0.09. Right: Wave DSI; p = 0.046. All statistical tests here are unpaired t-tests.

**Supplementary Fig. 6.**
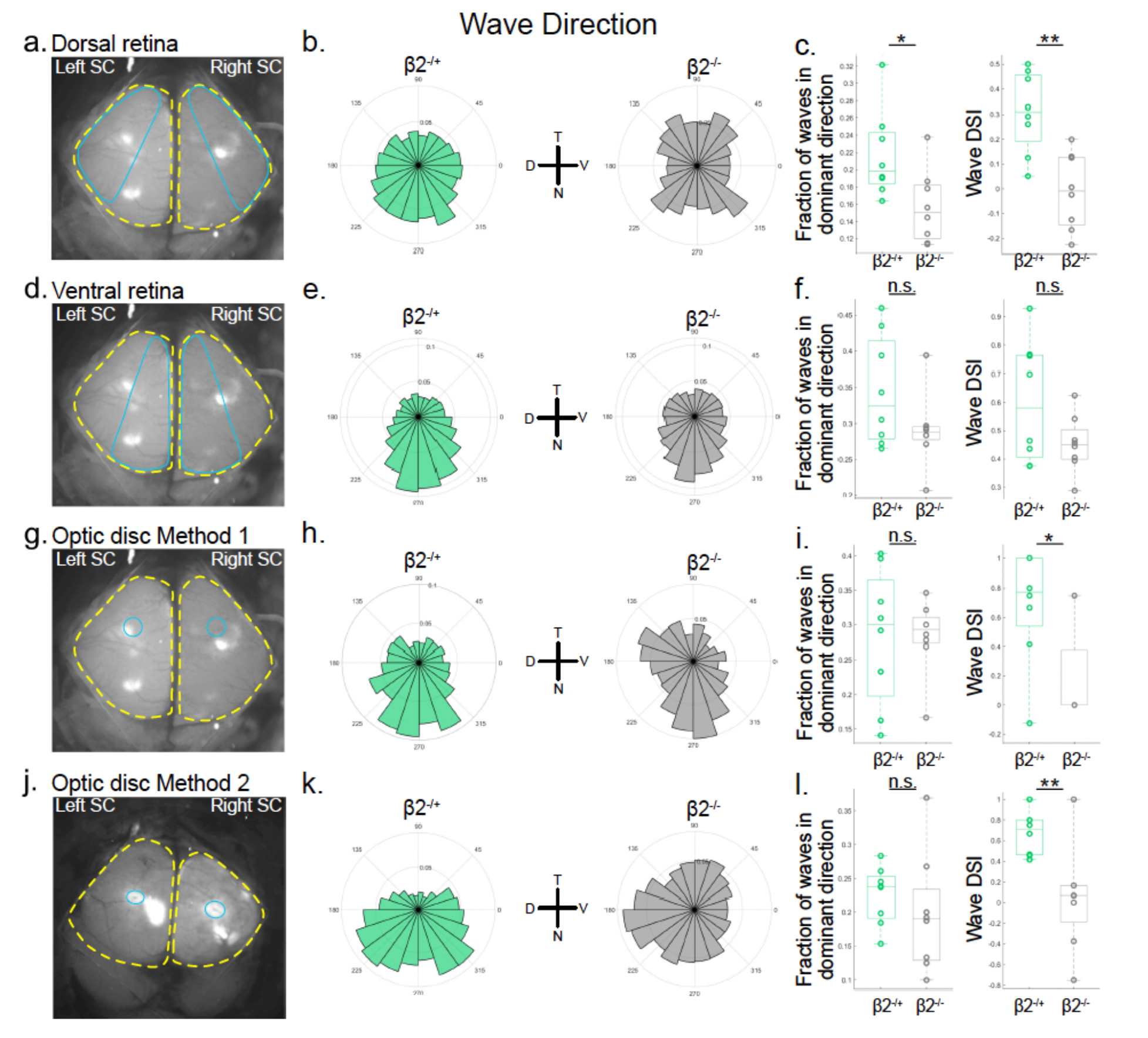
Restriction of SC field of view (FOV) to match retinal FOVs (related to Fig. 3). **a**. Example bird’s eye view of the SC during widefield one-photon calcium imaging. Yellow dashed lines indicate the outline of each hemisphere of the SC. Blue solid line indicates the restricted FOV used for subsequent analyses. Restricted SC FOV was chosen to be retinotopically matched with the dorsal retina. **b**. Frequency distribution of propagation directions of waves detected in the retinotopically matched SC area corresponding to dorsal retina in n = 8 β2^−/+^ mice and n = 8 β2^−/−^ mice (same data collected for Figure 3). **c**. Left: Proportion of waves traveling within 60° around the dominant temporal-to-nasal direction. Each data point represents one animal; p = 0.02. Right: Wave DSI; p = 0.0013. **d**. Restricted SC FOV was chosen to be retinotopically matched with the ventral retina. **e**. Frequency distribution of propagation directions of waves detected in the retinotopically matched SC area corresponding to ventral retina. **f**. Left: Proportion of waves traveling within 60° around the dominant temporal-to-nasal direction; p = 0.11. Right: Wave DSI; p = 0.1. **g**. Restricted SC FOV was chosen using previously reported retrograde fluorescent tracing experiments to correspond to the area adjacent to the optic nerve head in the retina (see Methods for details). **h**. Frequency distribution of propagation directions of waves detected within a SC FOV corresponding to the area adjacent to the optic nerve head in the retina. **i**. Left: Proportion of waves traveling within 60° around the dominant temporal-to-nasal direction; p = 0.99. Right: Wave DSI; p = 0.02. **j**. Restricted SC FOV was chosen using the patch of increased fluorescence expression in the SC (see Methods for details). **k**. Frequency distribution of propagation directions of waves detected within a SC FOV corresponding to the area adjacent to the optic nerve head in the retina. **l**. Left: Proportion of waves traveling within 60° around the dominant temporal-to-nasal direction; p = 0.42. Right: Wave DSI; p = 0.006. All statistical tests here are unpaired t-tests.

**Supplementary Fig. 7.**
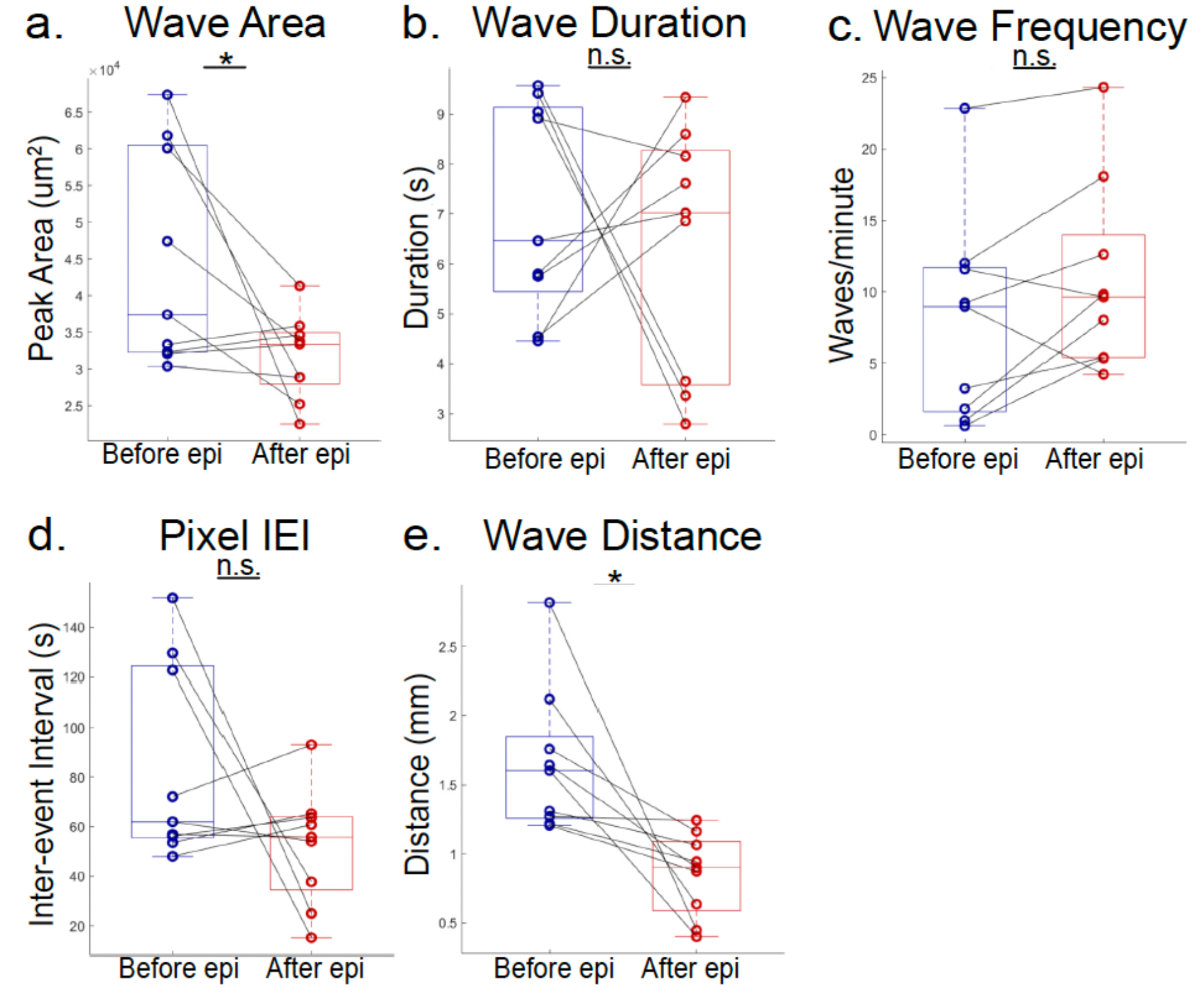
Other spatiotemporal properties after epibatidine injection (related to Fig. 4). **a**. Peak area of detected waves before and after an acute intraocular injection of 100 μM epibatidine. Each data point represents one animal; lines connect data from the same animal; n = 9 animals; p = 0.047. **b**. Median duration of detected waves; p = 0.62. **c**. Number of waves detected per minute; p = 0.07. **d**. Inter-event interval of activity detected at each pixel; p = 0.15. **e**. Wave propagation distance; p = 0.01. All statistical tests here are paired t-tests.

**Supplementary Fig. 8.**
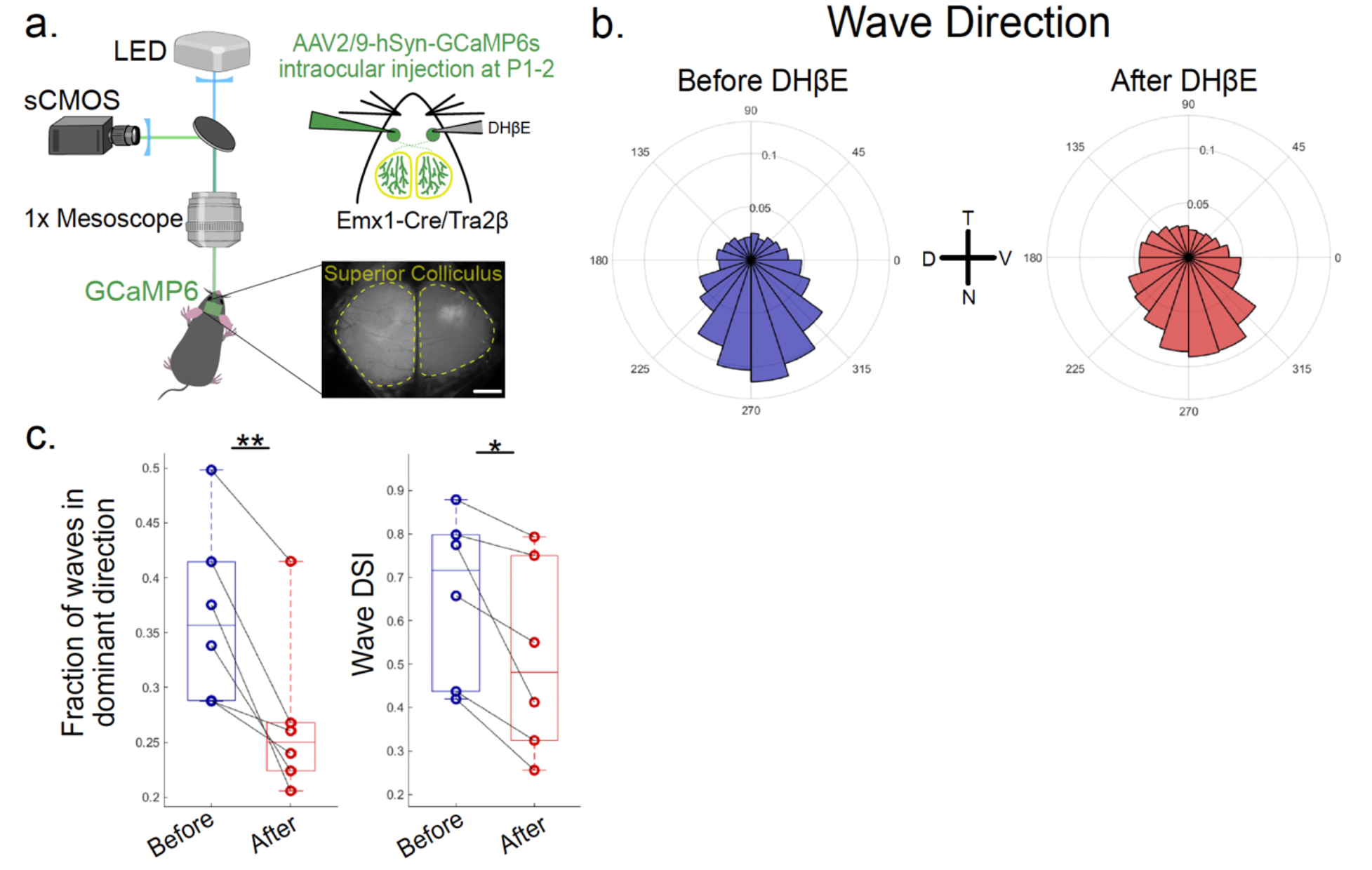
Stage III wave directionality is disrupted after DHβE injection (related to Fig. 4). **a**. Schematic of widefield single-photon microscopy used to perform calcium imaging of retinal ganglion cell axon activity in the SC of P9-11 Emx1-Cre/Tra2β^f/f^ (cortex-less mice). Spontaneous activity was recorded before and after an acute intraocular injection of DHβE or saline. Scale bar, 500 μm. **b**. Frequency distribution of propagation directions of waves before and after an acute injection of 200 μM DHβE; n = 6 animals. **c**. Left: Proportion of waves traveling within 60° around the dominant temporal-to-nasal direction. Each data point represents one animal; lines connect data from the same animal; p = 0.007. Right: Wave DSI; p = 0.024. Paired t-test.

